# The Organization of Serotonergic Fibers in the Pacific Angelshark Brain: Neuroanatomical and Supercomputing Analyses

**DOI:** 10.1101/2025.03.28.646017

**Authors:** Skirmantas Janušonis, Ralf Metzler, Thomas Vojta

## Abstract

Serotonergic axons (fibers) are a universal feature of all vertebrate brains. They form meshworks, typically quantified with regional density measurements, and appear to support neuroplasticity. The self-organization of this system remains poorly understood, partly because of the strong stochasticity of individual fiber trajectories. In an extension to our previous analyses of the mouse brain, serotonergic fibers were investigated in the brain of the Pacific angelshark (*Squatina californica*), a representative of a unique (ray-like) lineage of the squalomorph sharks. First, the fundamental cytoarchitecture of the angelshark brain was examined, including the expression of ionized calcium binding adaptor molecule 1 (Iba1, AIF-1) and the mesencephalic trigeminal nucleus. Second, serotonergic fibers were visualized with immunohistochemistry, which showed that fibers in the forebrain have the tendency to move toward the dorsal pallium and also accumulate at higher densities at pial borders. Third, a population of serotonergic fibers was modeled inside a digital model of the angelshark brain by using a supercomputing simulation. The simulated fibers were defined as sample paths of fractional Brownian motion (FBM), a continuous-time stochastic process. The results reproduced key features of serotonergic fiber densities in the telencephalon, a brain division with a considerable physical uniformity and no major “obstacles” (dense axon tracts). The study provides further evidence that serotonergic fibers can be successfully modeled as paths of a rigorously-defined stochastic process, and that a rich repertoire of self-organizing behaviors can be produced by axons that are inherently stochastic but also respond to external forces.

## 1. INTRODUCTION

Serotonergic axons (fibers) have been found in all studied vertebrate brains, where they form dense meshworks in many brain regions. Their ubiquity and morphological similarity across distant phylogenetic branches (cartilaginous and bony fishes, amphibians, sauropsids, mammals) suggests that they are an obligatory component of neural tissue, independent of clade-specific cytoarchitectonic specializations (e.g., the mammalian cerebral cortex) (Parent, 1981; Lillesaar, 2011; Yip et al., 2020; Awasthi et al., 2021; Bhat and Ganesh, 2023; Fujita et al., 2023; Biradar and Ganesh, 2024). Serotonergic fibers store and release serotonin (5-hydroxytryptamine, 5-HT) that can be detected by a large class of receptors (Bockaert et al., 2006), through volume or wiring neurotransmission (Papadopoulos et al., 1987; Smiley and Goldman-Rakic, 1996; Gianni and Pasqualetti, 2023). Some evidence exists that serotonin can directly modulate the membrane lipid bilayer (Dey et al., 2021) and can even enter cells by crossing their membrane (Andrews et al., 2022). Fundamentally, serotonergic signaling appears to support neuroplasticity (Lesch and Waider, 2012; Teissier et al., 2017; Cazettes et al., 2021; Ogelman et al., 2024; Page et al., 2024). No major neuroanatomical or electrophysiological defects have been detected in mice that cannot synthesize serotonin in the central nervous system (CNS), but these mice do show innervation and behavioral alterations (Migliarini et al., 2013; Montalbano et al., 2015; Mosienko et al., 2015). Notably, serotonergic fibers have no analogs in current artificial neural network architectures (Lee et al., 2022), where the loss of plasticity in deep continual learning remains a challenging problem (Dohare et al., 2024).

Recent studies have revealed a great diversity of transcriptional programs in the mouse serotonergic neurons (Okaty et al., 2019; Ren et al., 2019; Okaty et al., 2020) and also have shown that serotonergic fibers can regenerate in adult mammalian brains, with new trajectories (Hawthorne et al., 2011; Jin et al., 2016; Kajstura et al., 2018; Cooke et al., 2022). Most serotonergic axons appear to be unmyelinated, but myelinated serotonergic fibers have been reported in rodents and primates (Azmitia and Gannon, 1983; Westlund et al., 1992).

The processes that assemble serotonergic fibers into brain region-specific densities remain poorly understood. In mammals, the serotonergic fibers originate in the brainstem raphe nuclei and as a population reach all brain regions (Jacobs and Azmitia, 1992; Hornung, 2003). Different raphe nuclei and their subdivisions appear to preferentially target specific brain regions, but these projections are not fully segregated (Vertes, 1991; Vertes et al., 1999; Ren et al., 2018; Ren et al., 2019). In embryonic development, serotonergic fibers form some well-defined fascicles that carry fibers to the forebrain (Lidov and Molliver, 1982; Wallace and Lauder, 1983; Aitken and Tork, 1988; Slaten et al., 2010). However, these fascicles tend to travel along existing axon tracts produced by other, non-serotonergic axons (e.g., the medial forebrain bundle, the fasciculus retroflexus), that may serve only as contact guides (Lidov and Molliver, 1982; Wallace and Lauder, 1983; Jacobs and Azmitia, 1992). Growing serotonergic axons form specific adhesion structures on other neurites *in vitro* (Hingorani et al., 2022). A number of factors have been implicated in the growth, distribution, and survival of serotonergic fibers: the transcription factors Lmx1b and Pet-1 (Donovan et al., 2019; Kitt et al., 2022), protocadherin-αC2 (Chen et al., 2017; Katori et al., 2017), neurexins (Cheung et al., 2023), neuritin (Shimada et al., 2024), S100β (Azmitia and Whitaker-Azmitia, 1991; Whitaker-Azmitia, 2001), the brain-derived neurotrophic factor (BDNF) (Mamounas et al., 1995), serotonin itself (Migliarini et al., 2013; Nazzi et al., 2019; Nazzi et al., 2024), and others (Kiyasova and Gaspar, 2011). However, the relative roles of these factors (e.g., essential, supportive, injury-related), as well as their interactions, require further research. For example, transgenic mice lacking S100β appear to have no major defects in the development of serotonergic neurons (Nishiyama et al., 2002), and questions remain about serotonin levels required to significantly alter serotonergic fiber densities (Migliarini et al., 2013; Donovan et al., 2019).

We have recently introduced a novel approach that treats serotonergic fibers as paths of spatial stochastic processes, with no guiding cues (Janušonis and Detering, 2019; Janušonis et al., 2020). Such processes are random walk-like, but they can evolve in continuous time and have a complex autocorrelative structure. In particular, the trajectories of serotonergic fibers can be modeled as sample paths of fractional Brownian motion (FBM), a major generalization of normal Brownian motion that allows correlations between steps and exhibits long-range dependence (Mandelbrot and Van Ness, 1968; Biagini et al., 2010). Reflected FBM, a further extension necessary for computer simulations in bounded shapes (such as a brain), has been introduced only recently (Wada and Vojta, 2018; Guggenberger et al., 2019; Vojta et al., 2020). We have shown that FBM-fibers in a three-dimensional model of the mouse brain can automatically reproduce some regional serotonergic fiber densities described by neuroanatomical studies (Janušonis et al., 2023).

The proposed stochastic models are predictive in that they seek to simulate regional serotonergic fiber densities in any vertebrate brains – extant, extinct, or hypothetical. The results obtained in the mouse brain are promising (Janušonis et al., 2020; Janušonis et al., 2023), but they require validation in vertebrate brains that have different brain shapes and express different genes. Diverse brain shapes are important because reflected FBM is sensitive to the geometry of the bounding borders (Janušonis et al., 2020; Vojta et al., 2020). Since FBM has “memory,” this effect is not fully local and reflects the history of the fiber trajectory. Diverse genomes are important because the model assumes that the key properties of regional fiber densities can be predicted without detailed knowledge of axon guidance mechanisms. It should be noted that the model does not rule out such mechanisms and can include them as additional “forces” (e.g., modeled by “effective potentials” (Rizzo et al., 2013)) to further refine predictions. However, these extensions require quantitative definitions of such “forces” in the brain space, which are difficult to obtain from current experimental data.

Fish brains meet these two criteria. In particular, the extant cartilaginous fishes (sharks, skates, rays) produce a remarkable diversity of brain shapes (Yopak, 2012) and nearly maximize the phylogenetic distance to the mammals. In addition, the distributions of neurons in shark telencephala are much more uniform compared to those in mammalian telencephala which contain many differentiated cortical and subcortical structures. Therefore, a shark telencephalon can serve as a natural model of neural tissue that possesses a considerable degree of physical homogeneity, up to the natural brain borders. In computer simulations, these borders (the pial and ependymal surfaces) can be unambiguously modeled as impenetrable boundaries. Such biomechanical homogeneity does not imply chemoarchitectonic or functional homogeneities (unrealistic for any functional brain), but it eliminates the problem of complex physical “obstacles” (e.g., dense anatomical nuclei or axon tracts). These “obstacles” may differ in their permeability to serotonergic fibers (Janušonis et al., 2020) and viscoelastic properties (Chaudhuri et al., 2020; Antonovaite et al., 2021). In computer simulations, they introduce uncertainties and require careful treatment (Janušonis et al., 2020; Janušonis et al., 2023). It should be noted that nearly perfectly uniform environments can be created in cell cultures (Hingorani et al., 2022); however, these systems differ strongly from the natural brain environment, physically and chemically. In particular, they are usually two-dimensional, which has a major impact on the normal behavior of growing axons (Santos et al., 2020).

The extant sharks comprise around 500 species (Compagno et al., 2005) that can be divided into two superorders, the squalomorph sharks (Squalomorphii) and the galeomorh sharks (Galeomorphii; this group includes the great white shark). Relative to body size, the squalomorph sharks tend to have smaller brains (Striedter and Northcutt, 2020), with fewer differentiated cell masses (Butler and Hodos, 2005), which makes them convenient for the purpose of this study. Among several squalomorph species readily available along the coast of Southern California, we selected the Pacific angelshark *(Squatina californica)*, a representative of the angelshark order (Squatiniformes).

The angelshark body is superficially similar to that of rays; paleontological evidence suggests that this lineage dates back to the Late Jurassic (López-Romero et al., 2020; Maisey et al., 2020). Historically, angelsharks have been called “angels,” “monks,” and “bishops,” with colorful descriptions dating back to at least the 16^th^ century (McCormick et al., 1963; Compagno et al., 2005). The Pacific angelshark is found along the continental shelf of the east Pacific, typically on sandy flats (Compagno et al., 2005). To our knowledge, angelshark brains have been studied only at the gross anatomical level (Striedter and Northcutt, 2020), with no published reports on their cytoarchitecture or chemoarchitecture.

In this report, we first describe the basic neuroanatomy of the Pacific angelshark brain and visualize its serotonergic fibers. We then build its three-dimensional model, populate it with growing fibers modeled as reflected FBM paths, and use a supercomputing simulation to generate regional fiber densities. Finally, we compare these simulated densities with the biological fiber densities in the angelshark brain.

## 2. METHODS

### 2.1. Animals and Brain Preparation

Three adult Pacific angelshark *(Squatina californica)* specimens (two females, one male) were collected in 2017 from the UCSB Parasitology laboratory, with a post-mortem interval of 0-2 hours (after euthanasia with MS-222). The brains of the specimens were immediately removed, rinsed in 0.1 M phosphate-buffered saline (PBS, pH 7.2), and immersion-fixed in phosphate-buffered 4% paraformaldehyde at 4°C overnight. The fixed brains were immersed in 30% sucrose in PBS at 4°C for 2 days, transferred to a cryoprotectant solution (30% sucrose, 1% (w/v) polyvinylpyrrolidone (PVP-40), and 30% ethylene glycol in PBS) at 4°C for 2 days, and then moved (in cryoprotectant) to –20°C for indefinite storage. All procedures have been approved by the UCSB Institutional Animal Care and Use Committee.

### 2.2. Brain Sectioning

The brains stored in cryoprotectant were moved from –20°C to 4°C for one day and then transferred to 30% sucrose in PBS at 4°C for 2 days. They were divided into two rostro-caudal pieces, each of which was embedded in 20% gelatin (type A; Thermo Scientific #61199-5000) in a Peel-Away mold. An insect pin was pushed through the mold in the rostro-caudal orientation over the dorsal surface of the brain for physical support and as a fiducial marker for further alignment (Janušonis et al., 2023). After one hour at 4°C, the gelatin blocks were removed, trimmed, and immersed for 3 hours in undiluted formalin with 20% sucrose at room temperature. They were sectioned coronally from the rostral pole through the rostral myelencephalon at 40 µm thickness on a freezing microtome into 96-well trays with PBS. In order to avoid distance distortions in the rostro-caudal axis, damaged or missing sections were marked with empty wells. The processing of the two blocks was staggered to ensure the same exposure times.

### 2.3. The Serial Section Set

A subset of the complete section series from one female brain was used to visualize cell bodies and major fiber tracts. The same set was used to build a 3D-model of the Pacific angelshark brain for supercomputing simulations (section 2.7).

One-fourth of all ordered sections (with a constant rostro-caudal step of 160 µm) were mounted onto gelatin/chromium-subbed glass slides and allowed to air-dry. They were used to acquire three sets of images. First, the sections were imaged uncoverslipped in bright-field with a 1× objective. In these images, neural tissue is uniformly dark, which supports efficient capture of outer and inner (e.g., ventricle) brain contours. Next, the sections were briefly dipped in water and, while wet, were again imaged uncoverslipped with the same 1× objective. It revealed major fiber tracts that appeared dark against the remaining light tissue (due to light refraction). Finally, the same sections were Nissl-stained to reveal the basic cytoarchitecture of the neural tissue. They were rehydrated, stained with 0.25% thionine acetate (Millipore-Sigma #861340) for 13 sec, dehydrated in a graded series of ethanols, differentiated in 95% ethanol with 1.1% glacial acetic acid for 4 min, further dehydrated in absolute ethanol, cleared (defatted) in Xylenes, and coverslipped with Permount.

The remaining unmounted sections were moved to cryoprotectant at 4°C (with several changes over a few days) and eventually stored in 20 mL glass scintillation vials at –20°C until further processing.

### 2.4 Iba1 Immunohistochemistry

Some free-floating sections were stained for ionized calcium binding adaptor molecule (Iba1), also known as allograft inflammatory factor 1 (AIF-1) (all procedures were at room temperature unless otherwise indicated). Sections in cryoprotectant were allowed to slowly equilibrate to 4°C and thoroughly rinsed in PBS. They were blocked in 2% normal donkey serum (NDS; Jackson ImmunoResearch #017-000-121) for 30 min and incubated in rabbit anti-Iba1 IgG (1:500; FUJIFILM Wako Pure Chemical Corporation #019-19741) with 2% NDS and 0.3% Triton X-100 (TX) in PBS for two days at 4°C on a shaker. They were rinsed in PBS (three times, 10 min each), incubated in AlexaFluor 488-conjugated donkey anti-rabbit IgG (1:1000; ThermoFisher Scientific #A-21206) with 2% NDS in PBS for 90 min, rinsed in PBS (three times, 10 min each), mounted onto gelatin/chromium-subbed glass slides, allowed to air-dry in the dark, and coverslipped with the ProlongGold antifade mountant with the DNA stain DAPI (Thermo Fisher Scientific #P36931).

### 2.5 5-HT Immunohistochemistry

Sections at representative coronal levels were stained for 5-HT to visualize serotonergic fibers (all procedures were at room temperature unless otherwise indicated). Sections in cryoprotectant were allowed to slowly equilibrate to 4°C and thoroughly rinsed in PBS. They were blocked in 3% NDS for 30 min and incubated in goat anti anti-5-HT IgG (1:500; ImmunoStar #20079) with 2% NDS and 0.3% Triton X-100 (TX) in PBS for two days at 4°C on a shaker. They were rinsed in PBS (three times, 10 min each), incubated in Cy3-conjugated donkey anti-goat IgG (1:400; Jackson ImmunoResearch #705-165-147) with 2% NDS in PBS for 90 min, rinsed in PBS (three times, 10 min each), mounted onto gelatin/chromium-subbed glass slides, allowed to air-dry in the dark, and coverslipped with the ProlongGold antifade mountant with DAPI. Some sections were double-stained for 5-HT and Iba1. The procedure was the same, with the addition of the rabbit anti-Iba1 antibody and the AlexaFluor 488-conjugated donkey anti-rabbit antibody in the primary and secondary antibody steps (at the concentrations given in section 2.4).

### 2.6. Microscopy

Bright-field and epifluorescence imaging was performed on a Zeiss AxioImager Z1 system, using 1× (Plan-Neofluar, NA = 0.025), 5× (EC Plan-Neofluar, NA 0.16), and 10× (Plan-Apochromat, NA 0.45) objectives. Confocal imaging was performed on a Leica SP8 resonant scanning confocal system in three channels (Cy3, AlexaFluor 488, DAPI), using a 63×oil objective (NA 1.40) with the xy-resolution of 59 nm/pixel and the z-resolution of 299 nm/optical section. The figures show maximum-intensity projections.

### 2.7. The Brain Model

The images of the dry sections (separated by a constant rostro-caudal step of 160 µm; section 2.3) from the rostral pole to the mid-mesencephalon level were imported into the Reconstruct software (SynapseWeb) and aligned in the rostro-caudal axis, as described in our previous publications (Flood et al., 2012; Janušonis et al., 2023). Outer and inner brain contours were outlined in Photoshop by an expert neuroanatomist, imported into Wolfram Mathematica 14 (Wolfram Research, Inc.), and converted to point (*x*, *y*) arrays, as previously described (Janušonis et al., 2020; Janušonis et al., 2023).

Briefly, each closed contour was read as an ordered set of points, smoothed, bilaterally symmetrized, and transformed to an *N* × 3 matrix (where *N* is the number of the rows). The rows represented the consecutive *y*-coordinates (with no gaps), from the most dorsal level to the most ventral level of the contour. Each row contained three values: its *y*-coordinate and the leftmost and rightmost *x*-coordinates of the shape enclosed by the contour (at this *y*-coordinate). Both *x*- and *y*-coordinates were integers. Since this format cannot capture concavities oriented in the dorsoventral direction, they were coded as separate contours representing “forbidden” regions in the corresponding convex shape. In this two-dimensional (2D)-integer grid, the side of each square cell represented the physical distance of 12.5 µm in the physical brain.

The most caudal section (at the mid-mesencephalon level) was cloned 10 times to reduce distortions at the unnatural caudal border (the biological brain continues further and smoothly transitions to the spinal cord; the entire organ ends only at the caudal point of the spinal cord). This approach was justified by the relatively constant shape of the biological brain at this level (the large tectum and the tegmentum).

In the next step, a three-dimensional brain model was built from the section stack. Each two-dimensional section (with the virtual thickness of 160 µm) was subdivided into 13 thinner virtual sections (around 12.5 µm in thickness each) using linear interpolation. This produced a simulation grid with cells (voxels) that were cubes with sides corresponding to 12.5 µm in the physical brain.

### 2.8. The Supercomputing Simulation

The supercomputing simulation was performed as described in our previous study in the mouse brain (Janušonis et al., 2023), with some modifications.

The fiber densities were produced by 4800 growing fibers that were randomly seeded in the most caudal coronal section (representing the mid-mesencephalon). This decision was motivated by the tegmental location of the rostral raphe nuclei in all vertebrates (Butler and Hodos, 2005; Lillesaar, 2011) and by the presence of serotonergic neurons in the tectum of some cartilaginous fish species (Stuesse et al., 1991). Fibers were not seeded at more rostral levels (including the hypothalamus); however, relative fiber densities are insensitive to the location of the origin points after sufficiently long simulation times. Each fiber was represented by a three-dimensional path of a discrete, superdiffusive FBM (Qian, 2003): its trajectory moved according to the recursion relation *r*_*n*+1_ = *r*_*n*_ + ξ_*n*_, where *r*_*n*_ is the three-dimensional walker position, and the steps (increments) ξ_*n*_ are a three-component discrete fractional Gaussian noise. The statistically-independent *x*, *y*, and *z* components of ξ_*n*_ were Gaussian random numbers with mean zero and variance σ^2^. Each component had long-range correlations between its steps; the corresponding noise covariance function between steps *m* and *m* + *n* was given by 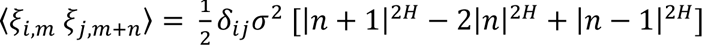, where *H* is the Hurst index, *δ_ij_* is the Kronecker delta, and *i*, *j* = *x*, *y*, *z* denotes the three space dimensions. The Fourier-filtering method was employed to generate these long-range correlated, stationary random numbers on the computer (Makse et al., 1996; Wada and Vojta, 2018; Vojta et al., 2020). The Hurst index was set at 0.8, based on our previous research in the mouse brain (Janušonis et al., 2020; Vojta et al., 2020; Janušonis et al., 2023), and the root mean-squared step size was set to *σ* = 0.2 grid units, corresponding to 2.5 µm in the physical brain (considerably less than the diameter of a single neuron). Each trajectory was composed of 2^25^ ≈ 33.6 million walk-steps. The length of the trajectories was sufficient for the relative densities to reach a steady state.

If a growing fiber encountered a boundary (i.e., an outer or inner contour), it was “reflected” by it (i.e., it was not allowed to cross it). Several reflection rules are available in simulations. They differ in computational complexity, but this choice has virtually no effect on simulation results, as we have shown previously (Vojta et al., 2020). In this simulation, a step that would push the leading fiber end into the forbidden region was simply not carried out. The entire sections at the most rostral and caudal levels were treated as reflecting boundaries.

After the simulation, the obtained densities were evaluated in non-overlapping cubes (composed of 2 × 2 × 2 grid cells, to suppress noise and achieve more robust estimates). The local density (*d*_s_) was determined by counting the total number of random-walk segments inside each cell. These local densities were normalized to the total sum of unity in the entire three-dimensional brain volume, to make the results independent of the arbitrarily chosen trajectory length. To facilitate comparisons between the simulated fiber densities and actual fiber densities in immunostained sections, the raw simulation densities were transformed to “optical densities” using the transformation *d*_o_ = exp (*kd_s_*) (Janušonis et al., 2020; Janušonis et al., 2023), with an empirically optimized *k* value (*k* = 10^8.2^). This transformation constrained density values to a finite interval (from zero to one), with lower (“darker”) values corresponding to higher fiber densities. Graphical density maps were produced in Wolfram Mathematica 14 using the built-in “GrayTones” color function.

All supercomputing simulations were written in Fortran 2018 and carried out on the Pegasus cluster at the Missouri University of Science & Technology, using parallel processing on several hundred CPU cores.

## 3. RESULTS

### 3.1. Anatomical Features of the Pacific Angelshark Brain

The Pacific angelshark (Fig. 1A) is a typical representative of the angelshark family (Squatinidae) and has a flattened body, dorsally-positioned eyes, and relatively large spiracles typical for bottom-dwelling shark species (Compagno et al., 2005). Its brain is relatively small compared to the body size (Fig. 1B, C), as expected for squalomorph sharks (Striedter and Northcutt, 2020). Considering that some of the major brain divisions (e.g., the telencephalon or cerebellum) can become greatly enlarged in various elasmobranch species (Hofmann and Northcutt, 2012; Yopak, 2012; Yopak et al., 2019; Striedter and Northcutt, 2020), none of these divisions appears “exaggerated” in the angelshark brain (Fig. 1B).

**Figure 1.**
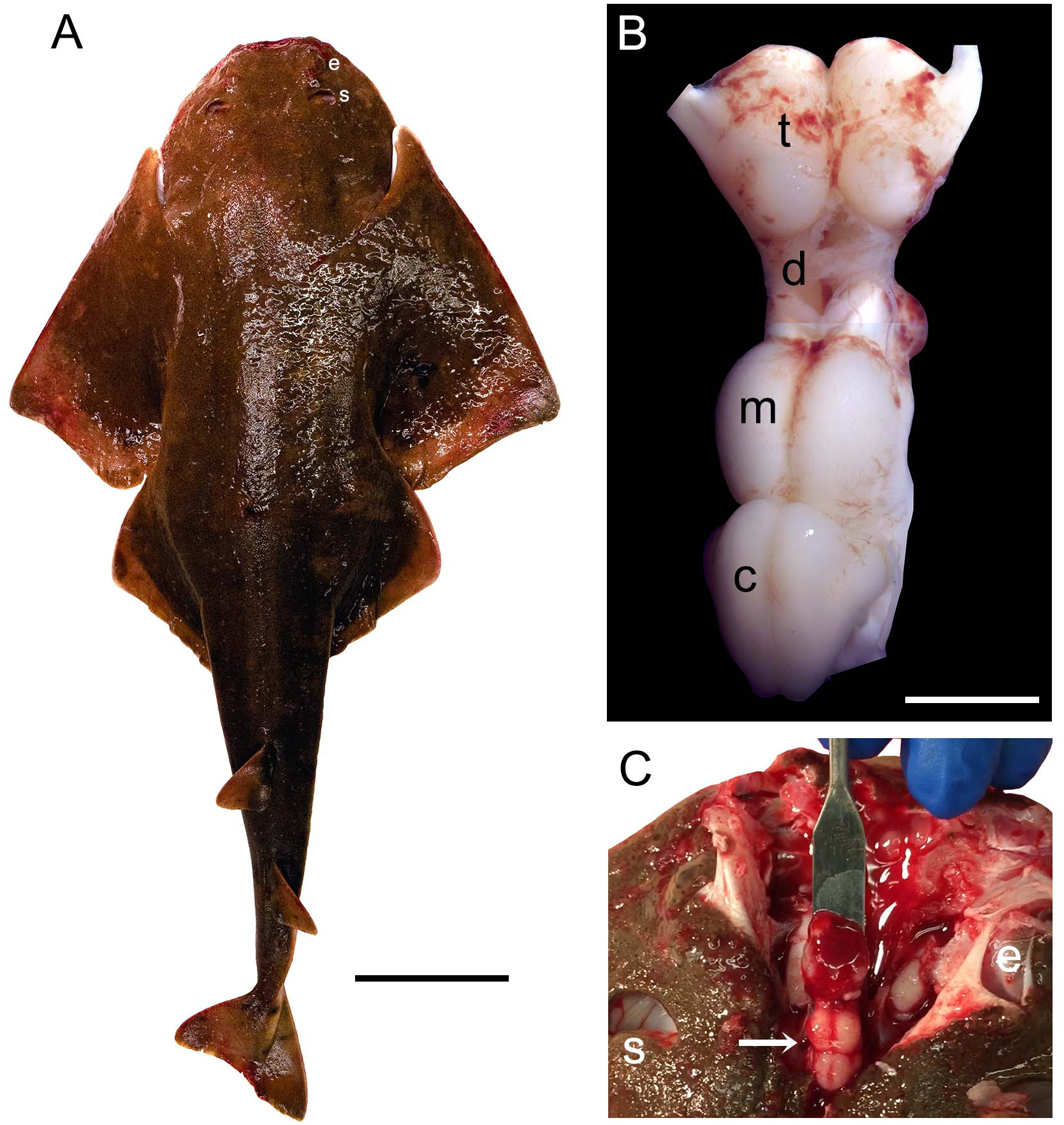
**(A)** The Pacific angelshark (a female specimen). Scale bar = 10 cm. (**B**) The Pacific angelshark brain (t, telencephalon; d, diencephalon; m, mesencephalon; c, cerebellum). The olfactory bulbs have been removed. Scale bar = 5 mm. (**C**) The Pacific angelshark brain (arrow) in situ. Abbreviations: e, eye; s, spiracle.

The basic neuroanatomy of the angelshark brain was captured in coronal sections that were imaged stained with a Nissl dye (to reveal cell bodies) and prior to the staining (to reveal highly refractive structures) (Fig. 2). The most prominent differentiated structure in the telencephalon was the area superficialis basalis (asb), a consistent feature in elasmobranch brains (Smeets et al., 1983; Smeets, 1998; Hofmann and Northcutt, 2012). The telencephalon had no strongly refractive structures, with the exception of the fasciculus basalis telencephali (fbt), the intensity of which became stronger as it approached the optic tract in diencephalon. In the diencephalon, extremely strong refraction was associated with the optic chiasm and the optic tract, confirming that the imaging correctly visualized major axon tracts. In stark contrast to the telencephalon, the entire mesencephalon and myelencephalon were strongly refractive, suggesting the presence of many densely packed axon tracts in this brain division. This observation was again consistent with fundamental anatomy, considering that cranial nerves III-XII, a universal input/output system in the vertebrate clade (Striedter and Northcutt, 2020), are directly associated with the brainstem.

**Figure 2.**
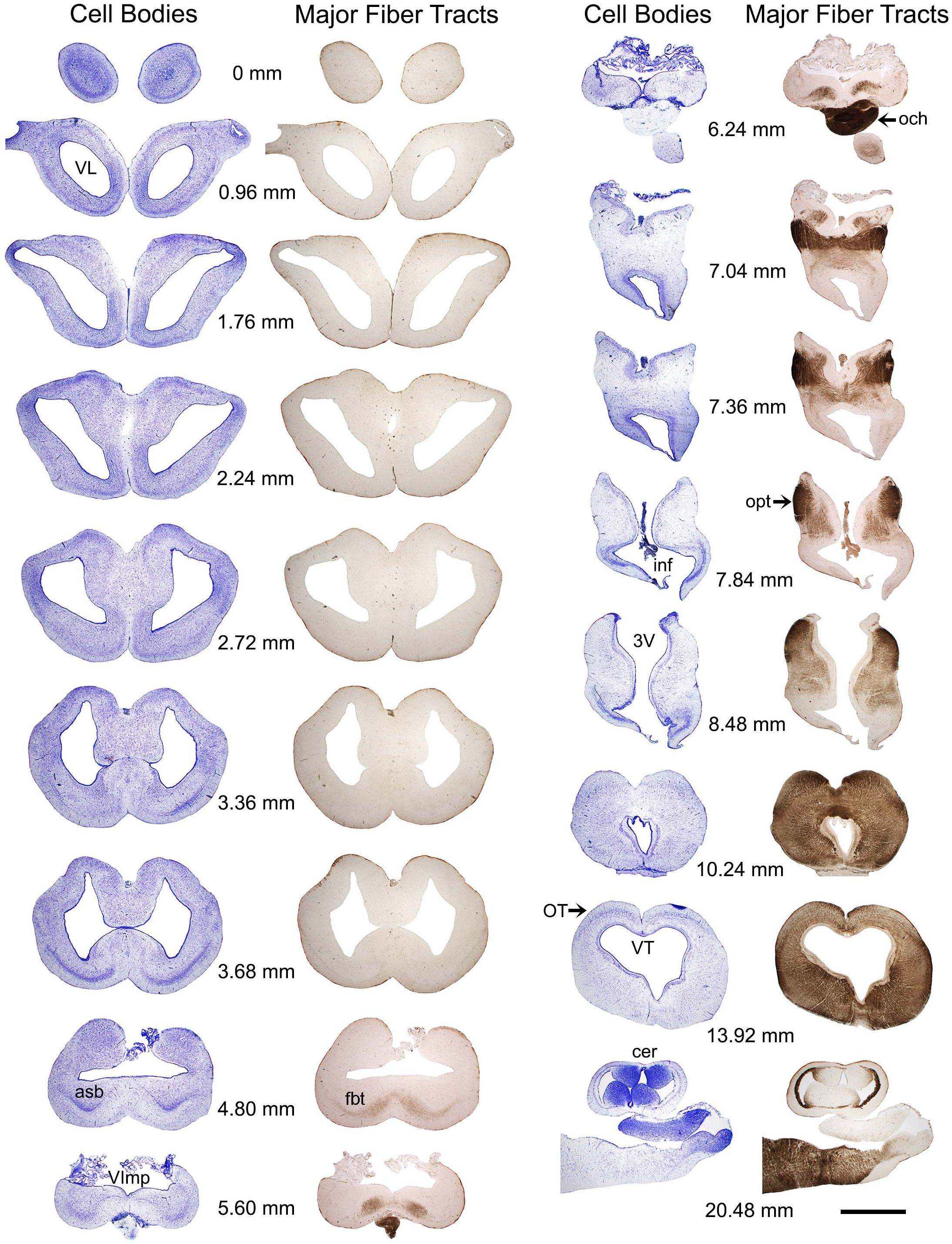
A subset of a serial set of coronal sections through a Pacific shark brain, from the rostral pole to the mid-cerebellum. The numbers represent the coronal distances (only key levels are shown; the rostro-caudal step is not constant). The sections were first imaged unstained, wet, and uncoverslipped to show the distribution of dense and highly refractive fiber tracts (the right columns) and then were stained with thionine to show the distribution of cell bodies (the left columns). At each coronal level, the sections in the right and left columns are the same. All sections were imaged in brightfield with a 1× objective. Abbreviations: 3V, third ventricle; asb, area superficialis basalis; cer, cerebellum; fbt, fasciculus basalis telencephali; inf, infundibulum; och, optic chiasm; opt, optic tract; OT, optic tectum; VImp, ventricle impar; VL, lateral ventricle; VT, tectal ventricle. Scale bar = 3 mm.

The highly diffractive structures were likely myelinated (de Bellard, 2016; Zalc, 2016; Verkhratsky et al., 2022), but the presence of myelin-associated proteins (e.g., myelin protein zero (MPZ)) was not directly examined in this study. Based on studies in other shark species, the angelshark telencephalon should contain several well-defined projections (Smeets, 1998; Hofmann and Northcutt, 2012). However, the majority of projections within the telencephalon are unmyelinated and loosely organized (Smeets et al., 1983; Smeets, 1998). The fbt is a notable exception and contains myelinated axons, as has been reported in another squalomorph species, the spiny dogfish (*Squalus acanthias*) (Smeets, 1998). In the angelshark, the fbt was located just ventral to the caudal asb (Fig. 3A-C). The asb contributes axons to the fbt (Hofmann and Northcutt, 2012), but probably not to the extent that was previously assumed (Smeets, 1998).

**Figure 3.**
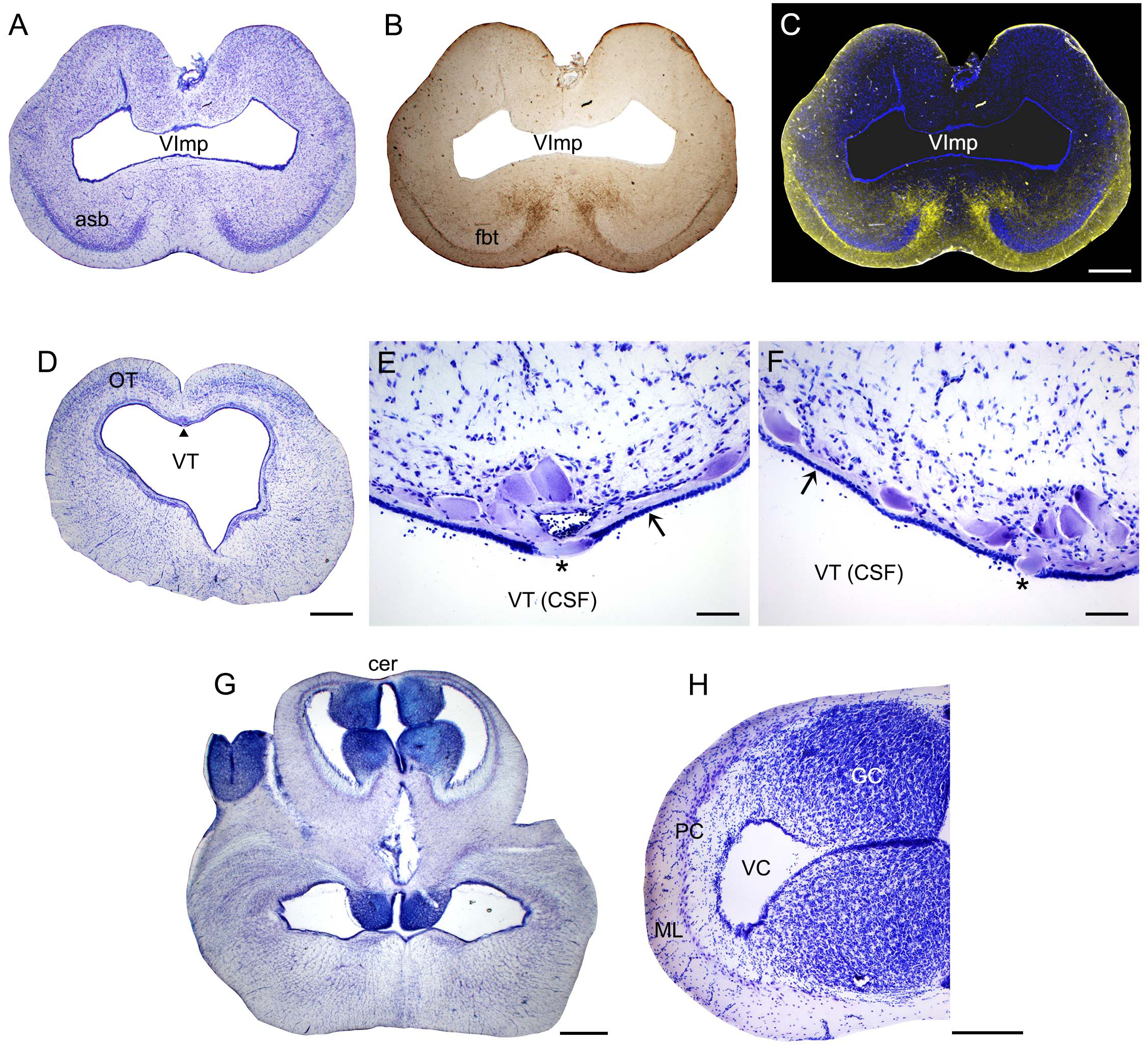
**(A)** A coronal section through the telencephalon stained to visualize cell bodies. (**B**) The same section before staining imaged wet to visualize major fiber tracts. (**C**) The two images superimposed (pseudocolored blue and yellow, respectively). (**D**) A coronal section through the mesencephalon stained to visualize cell bodies. The arrowhead indicates the mesencephalic trigeminal nucleus (Me5). (**E**, **F**) High-power images of Me5, showing thick subependymal processes (arrows) and cell bodies that cross the ependymal layer and come into direct contact with CSF (asterisks). (**G**) A coronal section through the cerebellum and myelencephalon stained to visualize cell bodies. (**H**) A high-power image of the cerebellum. Sections (A, D-H) were stained with thionine; all sections were imaged in brightfield. Abbreviations: asb, area superficialis basalis; cer, cerebellum; CSF, cerebrospinal fluid; GC, granule cells; fbt, fasciculus basalis telencephali; ML, molecular layer; OT, optic tectum; PC, Purkinje cells; VC, cerebellar ventricle; VImp, ventricle impar; VT, tectal ventricle. Scale bars = 1 mm (A-D, G), 100 µm (E, F), 400 µm (H).

A cluster of giant (around 100 µm in length) cells were present in the roof of the tectal ventricle (Fig. 3D). These cells corresponded to the mesencephalic trigeminal nucleus described in other shark species (Roberts and Witkovsky, 1975; Witkovsky and Roberts, 1975; 1976; MacDonnell, 1980; 1983; 1984; 1989). Consistent with these observations, the angelshark cells had thick subependymal processes, and some of their somata penetrated the ependyma to come into direct contact with the cerebrospinal fluid (CSF) (Fig. 3E, F). Somata inside the ependymal layer were consistently observed across the specimens, suggesting it was not a sectioning artifact.

The angelshark cerebellum had a structure typical for elasmobranchs (Fig. 3G, H). The granule cells were concentrated in the granular eminences (Butler and Hodos, 2005), long rostro-caudal columns that appeared nearly circular in some coronal sections.

We have previously shown that the ependymoglia of at least some galeomorph sharks shows strong immunoreactivity for ionized calcium binding adaptor molecule 1 (Iba1, AIF-1), a protein that is associated with immune responses and that is also a specific marker for the microglia in mammalian brains (Janušonis, 2018). Iba1 was also strongly expressed in the ependymoglia of the angelshark telencephalon (Fig. 4A), with ependymoglial processes often covering capillaries (Fig. B, C). Strong Iba1-immunoreactivity was also present in more caudal regions (Fig. D, E).

**Figure 4.**
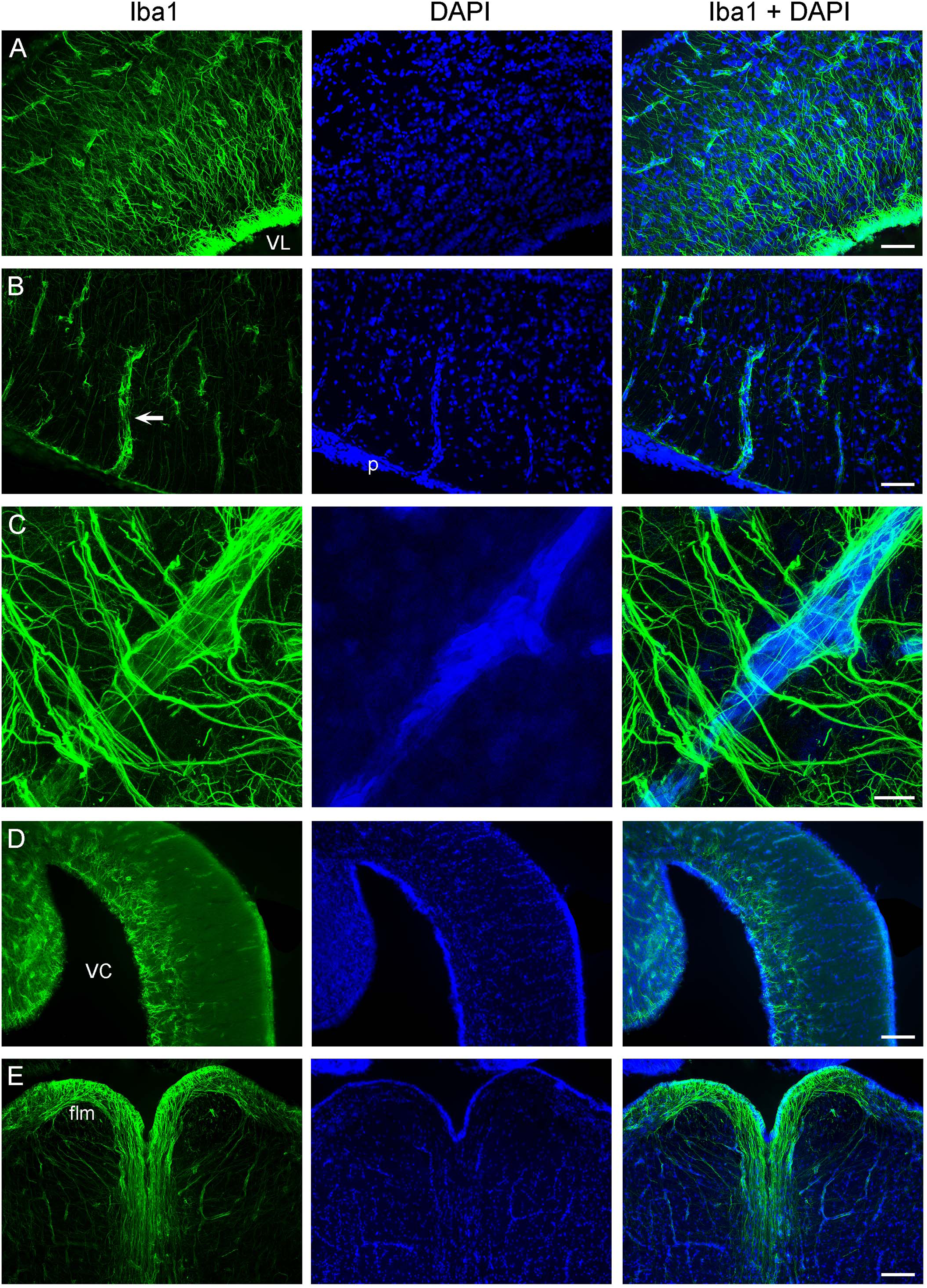
Iba1-immunoreactivity in the brain (cell nuclei are visualized with DAPI). (**A**) An epifluorescence image of Iba1+ ependymoglial cells in the telencephalon. The cell bodies are located at the lateral ventricle (VL) and extend their processes into brain parenchyma. (**B**) An epifluorescence image of a blood vessel (arrow) in the telencephalon. The blood vessel enters brain parenchyma through the pia (p) and is surrounded by Iba1+ processes. (**C**) A confocal image of a blood vessel in the telencephalon, with many fasciculated Iba1+ processes. The image is a maximum-intensity projection of 71 optical sections. (**D**) An epifluorescence image of Iba1-immunoreactivity in the cerebellum, at approximately the same coronal level as Fig. 3G (VC, cerebellar ventricle). (**E**) An epifluorescence image of Iba1+ fibers in the myelencephalon, at approximately the same coronal level as Fig. 3G (flm, fasciculus longitudinalis medialis). Scale bars = 100 µm (A, B), 25 µm (C), 200 µm (D, E).

### 3.2. The Anatomical Organization of Forebrain Serotonergic Fibers

Consistent with the general vertebrate pattern, a major source of serotonergic fibers in the angelshark brain is the brainstem raphe nuclei. Many of these fibers originate in the midbrain tegmentum (Fig. 5A) and spread through neural tissue. In addition, a major group of serotonin-immunoreactive cells was present in the angelshark hypothalamus (Fig. 5B). These cells appeared to make a major contribution to the forebrain serotonergic projections.

**Figure 5.**
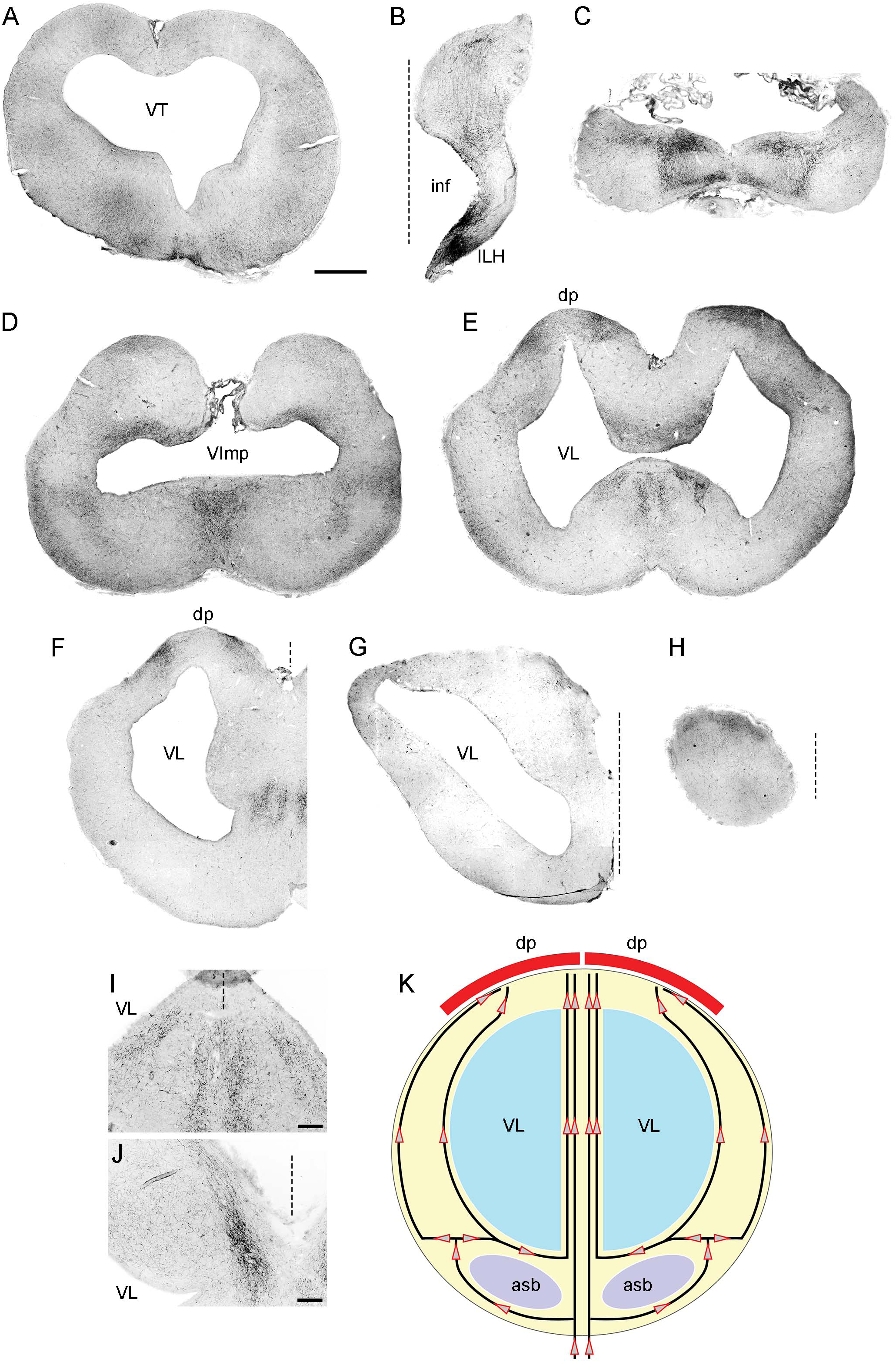
**(A-H)** Epifluorescence images of 5-HT-immunoreactivity at 8 coronal brain levels. Each image is a montage composed of 4-32 images captured with a 5× objective. The montages were converted to grayscale and digitally inverted (higher densities of serotonergic fibers are darker). The original montages (at full resolution and in color) are available in the Supplementary Material. (**I**) A higher-power image of 5-HT-immunoreactivity in a region corresponding to the ventromedial region of (E). Four streams of serotonergic fibers are apparent. (**J**) A higher-power image of 5-HT-immunoreactivity in a region corresponding to the dorsomedial region of (G). A medial stream of serotonergic fibers entering the dorsal pallium is apparent. (**K**) The hypothesized paths of serotonergic fibers. The key observations in (A-J) can be explained by assuming that the forebrain serotonergic fibers tend to accumulate close to pial and ependymal surfaces (perhaps due to their intrinsic stochasticity) but are additionally pulled by a deterministic force exerted by the dorsal pallium (dp). Abbreviations: asb, area superficialis basalis; dp, dorsal pallium; ILH, inferior lobe of the hypothalamus; inf, infundibulum; VImp, ventricle impar; VT, tectal ventricle. The midline is indicated by dashed lines. Scale bars = 1 mm (A-H), 200 µm (I, J).

The major serotonergic pathways in the forebrain were readily identifiable with 5-HT immunohistochemistry, revealing a relatively complex picture of transitions in the caudo-rostral axis (Fig. 5B-H). However, this sequence could be explained by a dynamic interplay of only two driving factors: (*i*) a tendency of fibers to accumulate at tissue borders and (*ii*) an attracting force generated by the dorsal pallium (Fig. 5K). This parsimonious approach facilitates the following description of the key pathways, but further confirmatory research is required.

In the diencephalon, many fibers emerged from strongly serotonin-immunoreactive cells that surrounded the ventricle in the inferior lobe of the hypothalamus (a large structure in shark brains). In this ventral region, fibers tended to accumulate at high densities near the ventricular and pial surfaces, with no identifiable orientation (Figs. 5B, 6A). In the dorsal region at the same coronal level, fibers formed a distinct oriented stream that coursed in the dorsal direction and then made a lateral turn to approach the habenula (Fig. 6A, B). Some of these fibers appeared to emerge from an initially disorganized cluster of fibers located midway between the ventral and dorsal regions (Fig. 6C). The fiber stream entered the habenula at the initial approach angle, but some fibers appeared to reorient themselves to next proceed to the caudal telencephalon (Fig. 6D).

**Figure 6.**
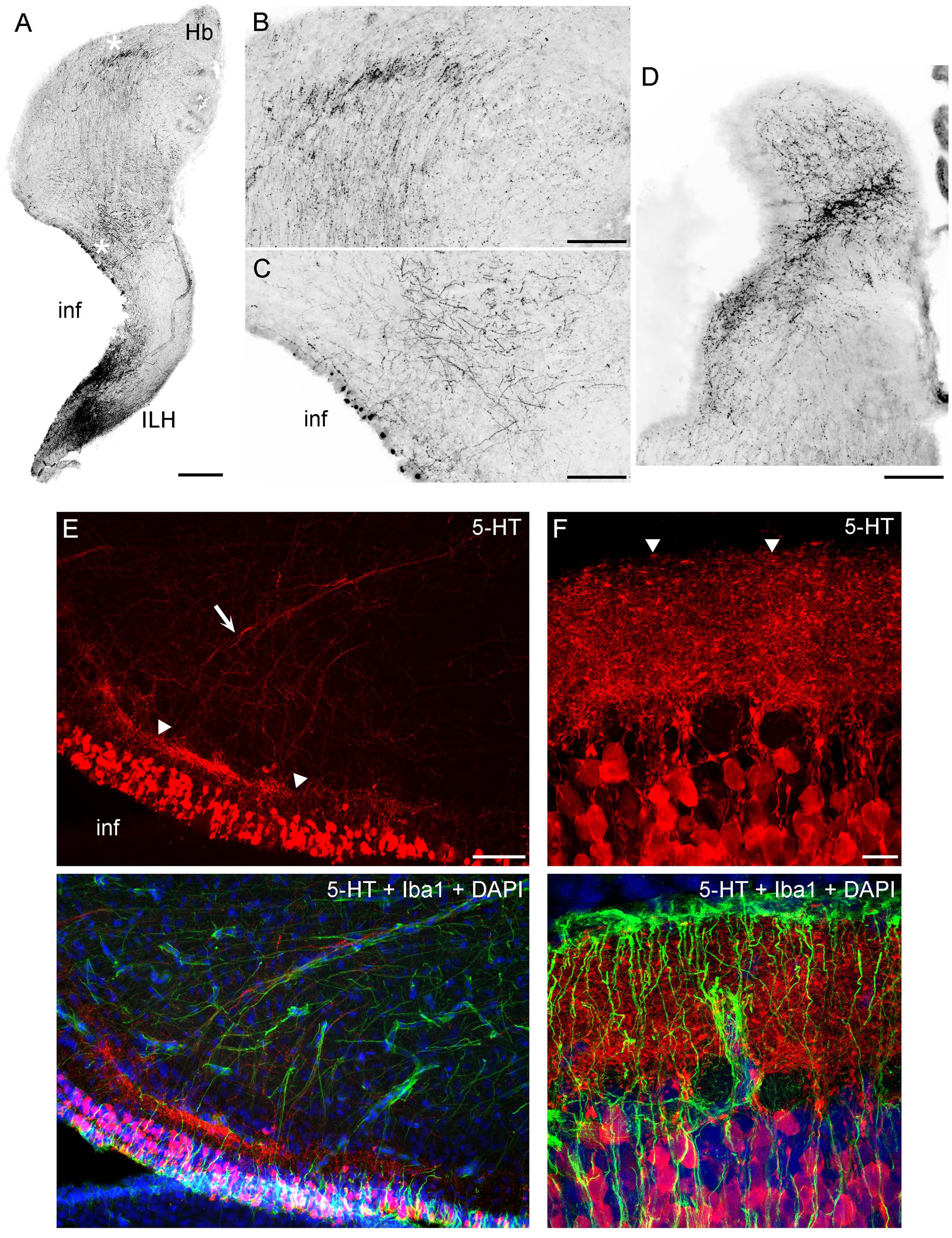
**(A-D)** Epifluorescence images of 5-HT immunoreactivity in coronal sections through the diencephalon. The images were converted to grayscale and digitally inverted (higher densities of serotonergic fibers are darker). (**A**) An image of one side of the diencephalon (corresponding to Fig. 5B). (**B**, **C**) Two enlarged regions of (A) (marked with asterisks). In (B), note the strongly oriented fibers coursing toward the habenula. (**D**) A stream of fibers in the habenula, at a coronal level adjacent to that of (A). (**E**) An epifluorescence image of the hypothalamus immunostained for 5-HT and Iba1 (with DAPI to visualize nuclei). (**F**) A confocal image of the caudal hypothalamus immunostained for 5-HT and Iba1 (with DAPI to visualize nuclei). The image is a maximum-intensity projection of 35 optical sections. Abbreviations: ILH, inferior lobe of the hypothalamus; inf, infundibulum; Hb, habenula. Scale bars = 500 µm (A), 200 µm (B-D), 100 µm (E), 20 µm (F).

In the caudal telencephalon, fibers again accumulated at the pial and ventricular surfaces, with an additional dorsal pull (Fig. 5C). Most fibers moved around the expected location of the fbt (Fig. 2 at 5.60 mm), consistent with the poor penetration of dense axon tracts by serotonergic fibers (Janušonis et al., 2020). More rostrally, fibers continued to accumulate near the pial and ventricular surfaces (Fig. 5D). In addition, a medial stream of fibers emerged at the coronal level of the ventricle impar. This stream could be explained by the opening of the median “land-bridge” between the ventral and dorsal parts of the telencephalon (the septal and medial pallial regions). This bridge splits the single ventricle impar (Fig. 5D) into the two lateral ventricles more rostrally (Fig. 5E, F) and can serve as a shortcut between the ventromedial telencephalon and the dorsal pallium. Consistent with this possibility, fibers formed four distinct medial streams, two on each side (Fig. 5F). Of the two streams, the more lateral one approached the bridge as a continuation of the fibers that spread near the ventricle (Fig. 5E, I). The other stream, adjacent to the median symmetry line, appeared to contain fibers that initially accumulated near the ventral pial surface but then were pulled in the dorsal direction (toward the medial pallium) when the more direct path to the dorsal telencephalon opened up (Fig. 5E, I, K).

In the mid-level telencephalon, two high-density clusters of fibers were present in the dorsal pallium (Fig. 5F). These clusters appeared to correspond to (*i*) fibers that approached the dorsal pallium moving around the ventricles (i.e., taking the longer lateral route), primarily staying close to the pial and ventricular surfaces, and (*ii*) fibers that used the shorter median route (Fig. 5K). The median projection to the dorsal pallium was directly visualized at a more rostral level (Fig. 5G, J). The most rostral pole of the brain still had a higher accumulation of fibers at the dorsal surface (Fig. 5H).

High-resolution imaging of the serotonin-containing cells in the periventricular hypothalamus (Fig. 6E-F) suggests a highly organized structure that bears some anatomical resemblance to the serotonergic system in the mammalian gut (enterochromaffin cells and serotonergic neurons in the gut wall) (Furness and Stebbing, 2018; Gershon and Margolis, 2021). These cells were uniform in their ovoid morphology (around 20 µm in length) and were densely packed around the ventricle in a continuous layer (around 100 µm thick). At some coronal levels, a distinct, dense serotonin-immunoreactive plexus was present under the cell-body layer, with fiber fascicles emerging from it (Fig. 6E). This plexus could be nearly as thick as the cell-body layer and appeared to contain extremely tightly packed neurites whose individual paths could not be distinguished in confocal microscopy at the limit of optical resolution (Fig. 6F). The fascicles moved dorsolaterally, entering the described pathway toward the habenula. However, this analysis cannot rule out that some of these fibers were contributed by more caudal regions (e.g., the raphe nuclei).

A systematic analysis of serotonergic fibers at the pial surface of the telencephalon revealed universally elevated densities (Fig. 7). Fiber densities showed some regional variation but were consistently higher near the pia with respect to the underlying (deeper) tissue.

**Figure 7.**
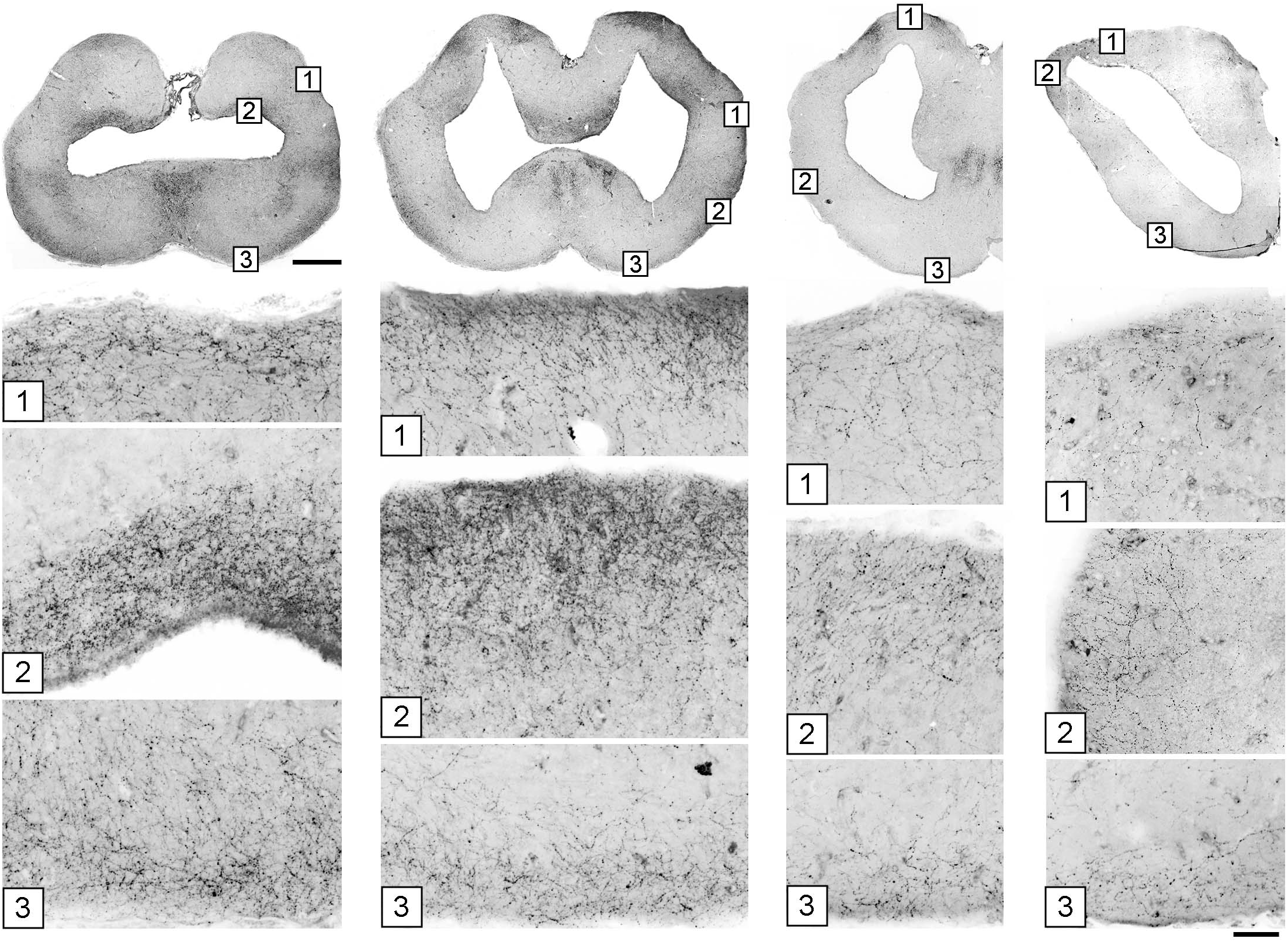
The accumulation of serotonergic fibers near tissue borders in the forebrain. The images in the top row correspond to Figures 5D-G. Under each coronal level, the high-power images show the accumulation of 5-HT+ fibers in border regions marked with corresponding numbers. The sections were immunostained for 5-HT, imaged in epifluorescence, converted to grayscale, and digitally inverted (higher densities of serotonergic fibers are darker). Some high-power images are rotated to orient the edge horizontally. Scale bars = 1 mm (top row), 100 µm (all panels below).

The angelshark brainstem poses challenges in computational simulations because it contains many dense axon tracts (“obstacles”) (Fig. 2); therefore, it was not a focus of this study. Serotonergic fibers in the brainstem are produced by the raphe nuclei that were well developed in the angelshark brain (Fig. 8A, B). Notably, serotonergic fibers did not produce a detectable accumulation at the pial and ventricular surfaces of the optic tectum (Fig. 8C-F).

**Figure 8.**
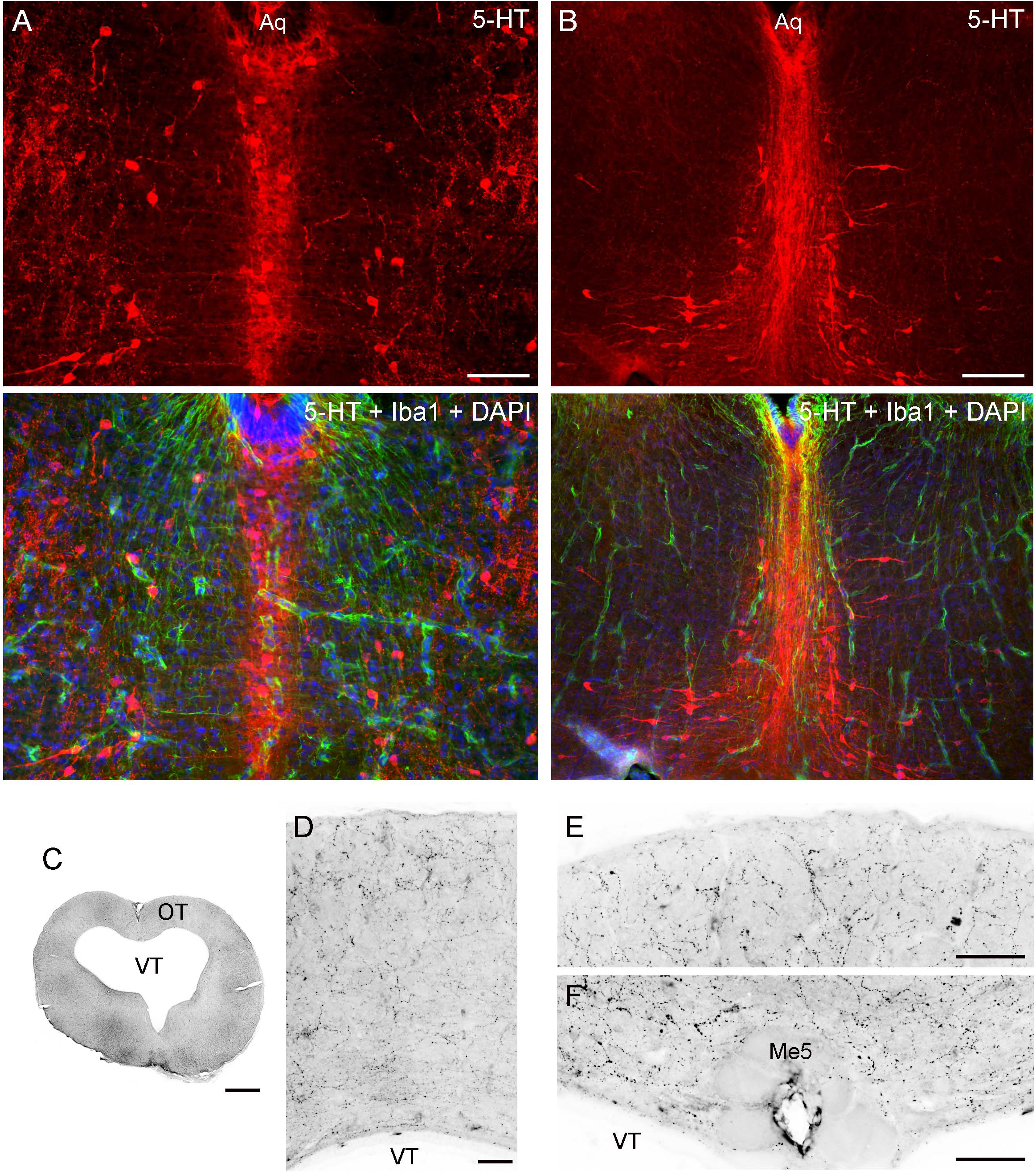
(**A, B**) Epifluorescence images of the raphe nuclei in coronal sections through the tegmentum stained for 5-HT and Iba1 (with DAPI to visualize nuclei). (**C**-**F**) Epifluorescence images of 5-HT immunoreactivity in coronal sections through the mesencephalon. The images were converted to grayscale and digitally inverted (higher densities of serotonergic fibers are darker). (**C**) An entire section corresponding to Figure 5A. (**D**) An enlarged image of the optic tectum (OT). (**E**) A high-power image of the superficial OT. (**F**) A high-power image of the periventricular OT. Abbreviations: Aq, aqueduct; Me5, mesencephalic trigeminal nucleus; VT, tectal ventricle. Scale bars = 100 µm (A, D-F), 200 µm (B), 1 mm (C).

### 3.3. The Supercomputing Simulation of Serotonergic Fibers

The described anatomical data was used to build and verify a predictive computational model of serotonergic fiber densities. The model was deliberately minimal in that it consisted only of fractional Brownian motion (FBM) trajectories, or idealized fibers, moving unguided within a three-dimensional shape that precisely matched the geometry of the Pacific angelshark brain. Despite its conceptual simplicity, the model was computationally expensive (due to the inherent mathematical properties of FBM) and required supercomputing resources. The hypothesized force toward the dorsal pallium was not included because the current biological information is insufficient to define it quantitatively.

An accurate, three-dimensional model of the Pacific angelshark brain was built based on an anatomical series of sections. The 2D-shapes at each coronal level closely followed the contours of physical sections and were minimally edited to only smooth the contours and achieve a perfect bilateral symmetry (Fig. 9). Next, 4800 fibers were launched from the brainstem and were allowed to roam in the brain shape with no internal obstacles. After the fibers achieved stable regional densities (Fig. 10), they were compared to those in the biological angelshark brain.

**Figure 9.**
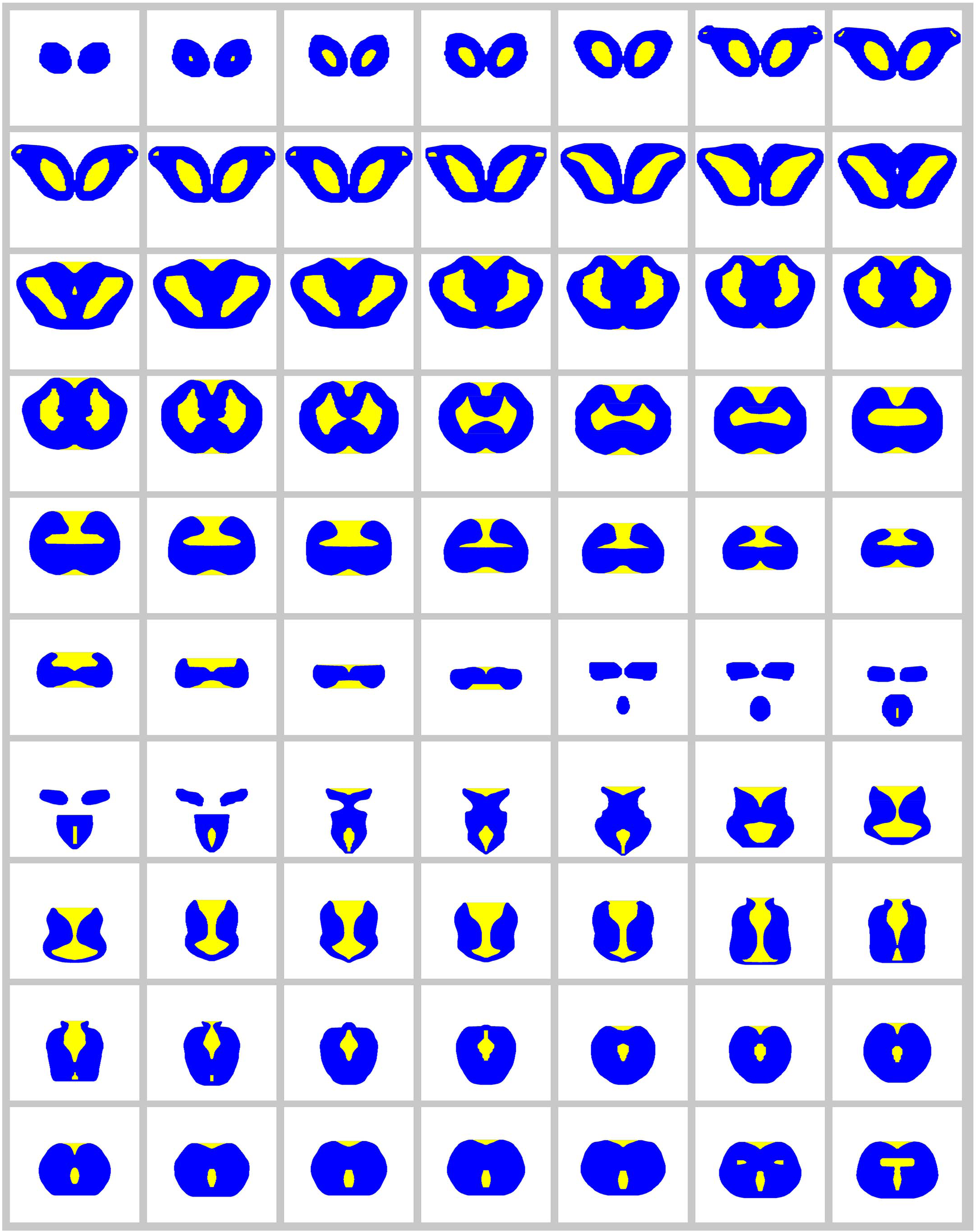
The complete series of digitized (and bilaterally symmetrized) images used to build a 3D-model of the angelshark brain for supercomputing simulations. The regions available for fibers are blue and the forbidden regions are yellow.

**Figure 10.**
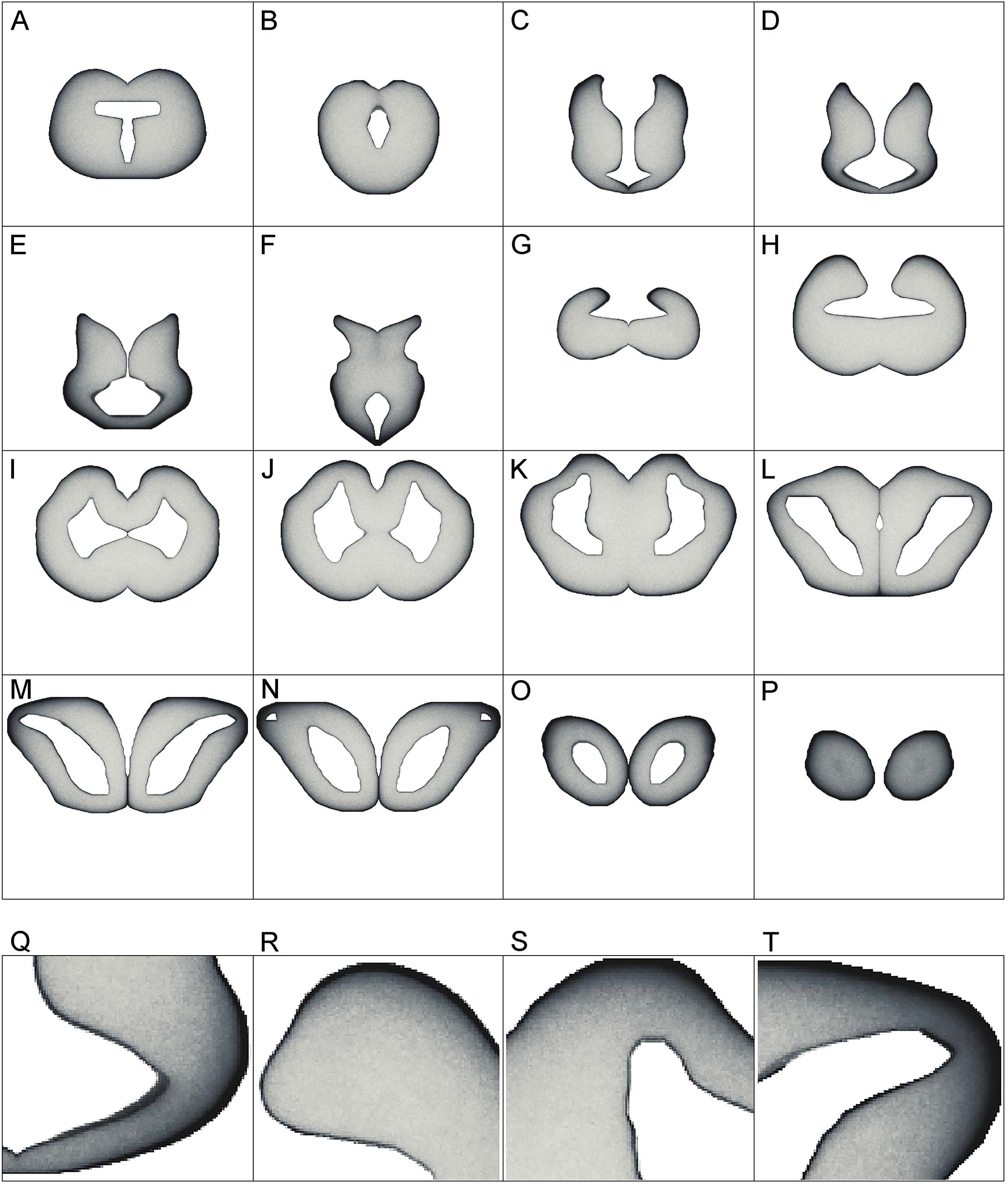
**(A-P)** The simulated densities of serotonergic fibers modeled as stochastic paths of fractional Brownian motion (FBM). Key coronal levels are shown. In the simulation, the fibers receive no guiding cues; regional density differences are the result of the brain geometry and the fundamental properties of FBM. Higher densities are darker, to facilitate comparisons with Figure 5. The images have no artificial contour lines (borders appear black because of fiber accumulation). (**Q**-**T**) Enlarged regions of (D), (H), (K), and (M), respectively.

Generally, fibers accumulated near tissue borders, especially at the pial border (Fig. 10A-P). This result was highly consistent with the elevated densities of serotonergic fibers at the pial border of the angelshark telencephalon (Fig. 7). In the diencephalon, a dense accumulation of simulated fibers was observed in the region surrounding the ventricle in the inferior lobe of the hypothalamus (Fig. 10D-F, Q). This pattern was similar to the elevated densities of serotonergic fibers in this region of the biological brain (Fig. 6A). The simulated densities were also high at the pial surface of the dorsal lobes overhanging the ventricle impar of the telencephalon (Fig. 10H, R), as well as in the prominent bulges of the dorsal pallium (Fig. 10K, S). This pattern was again consistent with the serotonergic fiber densities in the biological brain (Fig. 5D, E). Increased simulated and biological densities were present in the rostral dorsolateral telencephalon near the attachment of the stalk of the olfactory bulbs (Fig. 10M, N, T; Fig. 5G).

## 4. DISCUSSION

The gross morphology and basic cytoarchitecture of the Pacific angelshark brain share strong similarity with those of the spiny dogfish (*Squalus acanthias*), another squalomorph shark in a different (Squaliformes) order (Smeets et al., 1983; Smeets, 1998; Butler and Hodos, 2005). Compared to other sharks, where the cerebellar structures can make up over 40% of the brain (New, 2001; Montgomery et al., 2012), the angelshark cerebellum is small, probably because this species is a sedentary, ambush predator. A comprehensive rostro-caudal set of coronal sections was produced in this study (Fig. 2; see Data Availability), but we primarily focused on the brain geometry and the physical distribution of cell bodies (important for the modeling of serotonergic fibers). We did not investigate deeper levels of neural tissue organization that would require chemoarchitectonic or tract-tracing analyses (Smeets, 1998; Paxinos et al., 2022). Specifically, regions that appear homogeneous in the overall distribution of cells may contain well-defined subregions that are distinctly different in their functional network profiles (e.g., neurotransmitters, receptors, projections). With these limitations, this study contributes to comparative neuroanatomy, including two notable observations.

First, strong Iba1-immunoreactivity was observed in the ependymoglial (and potentially other) cells in the angelshark brain. This finding extends our previous report (Janušonis, 2018) and leads to the conclusion that, generally, Iba1 is expressed in the brains of both squalomorph and galeomorph sharks. However, it may not be expressed in some shark branches that diverged early (Janušonis, 2018). It is unknown whether the detected molecule is analogous to the mammalian Iba1, expressed in microglia (Wake et al., 2009), or is a functionally different molecule with a strong structural similarity to Iba1. Extensive research has shown that Iba1, also known as allograft inflammatory factor 1 (AIF-1), is highly conserved across the animal kingdom (Barca et al., 2017).

Second, a prominent mesencephalic trigeminal nucleus (Me5) was found in the angelshark brain. This nucleus, located on the midline in the roof of the tectal ventricle, contains gigantic cells (measuring 50-100 µm across) with thick processes that extend laterally just below the ependyma. Some cell bodies protrude ventrally into the ventricle and appear to reach the cerebrospinal fluid (CSF). This remarkable arrangement was studied in other shark species in a series of publications in the 1980s (MacDonnell, 1980; 1983; 1984; 1989), which followed earlier light microscopy, electron microcopy, and electrophysiological studies (Roberts and Witkovsky, 1975; Witkovsky and Roberts, 1975; 1976). This nucleus is interesting because it is the only primary sensory ganglion located within the CNS (Witkovsky and Roberts, 1975; Blumenfeld, 2022) and because of its role in clinical neurology (Blumenfeld, 2022). In the human brain, Me5 neurons form two lateral clusters and are known to mediate the monosynaptic jaw-jerk reflex. This reflex may be undetectable in healthy individuals but become unmasked in conditions such as amyotrophic lateral sclerosis (ALS) (Blumenfeld, 2022). A detailed electrophysiological study has described a similar reflex in the smallspotted catshark (*Scyliorhinus canicula*) and the dusky smoothhound (*Mustelus canis*) (galeomorph sharks, also known as dogfish), which suggests that this fast jaw-closing mechanism has been conserved in vertebrates (Roberts and Witkovsky, 1975). Our analysis supports an earlier hypothesis that, in sharks, Me5 activity (and the associated jaw closing) can be directly modulated by signals in the CSF (MacDonnell, 1980). This connection may have been lost in the reptiles and mammals, where Me5 is no longer adjacent to the ventricular wall (MacDonnell, 1980). We lastly note that the size of the Me5 neurons in the angelshark may be related to its powerful, extensible jaws that snap upward to capture prey within the striking distance of the immobile, camouflaged animal.

The angelshark telencephalon is relatively uniform in its spatial distribution of cell bodies (with the exception of the asb) and is virtually devoid of large-scale “obstacles” in the form of major axon tracts (with the exception of the fbt). It makes it a convenient natural model for large-scale simulations of serotonergic fibers. By comparison, a number of major axon tracts are present in all divisions of the adult mammalian brain, including the telencephalon (e.g., the anterior commissure, the corpus callosum). These tracts can be impenetrable to serotonergic fibers to various degrees (Lidov and Molliver, 1982; Janušonis et al., 2020). Mouse telencephalic regions also show regional differences in viscoleasticity, which are only partially understood (Antonovaite et al., 2021).

Compared to the telencephalon, the angelshark brainstem is highly refractive, likely due to densely packed axon tracts (Smeets, 1998). This observation is supported by viscoelastic measurements in the zebrafish (*Danio rerio*) brain that show that fish telencephalon is more viscous than the brainstem (which is stiffer) (Jordan et al., 2022). In the brainstem, axon tracts are expected to have different orientations and introduce strong local anisotropies. The current experimental data is insufficient to model these anisotropies accurately.

The primary sources of serotonergic fibers in the angelshark brain are the raphe nuclei of the brainstem and the inferior lobe of the hypothalamus. Raphe clusters of serotonergic neurons is a universal vertebrate feature (Jacobs and Azmitia, 1992; Stuesse et al., 1995; Hornung, 2003; Rodrigues et al., 2008; Lillesaar, 2011; Fonseca et al., 2021). Serotonergic hypothalamic neurons have been reported in other shark species, with additional clusters in other brain regions, such as the preoptic area, the pretectal area, and the habenula (Stuesse et al., 1991; Carrera et al., 2008; Lillesaar, 2011). However, serotonin immunoreactivity in these additional regions may be developmentally transient (Carrera et al., 2008), consistent with similar observations in the development of bony fish (Ebbesson et al., 1992). Hypothalamic serotonergic neurons also have been reported in reptiles, but not consistently across species (Rodrigues et al., 2008). They appear have been lost in mammals, with the only but remarkable exception of the monotremes (egg-laying mammals) (Manger et al., 2002).

Both raphe and hypothalamic neurons may contribute serotonergic fibers to the telencephalon. A developmental study of the smallspotted catshark has shown the presence of pioneering fibers in the telencephalon after the differentiation of the raphe serotonergic neurons but just before the sudden (“explosive”) appearance of the hypothalamic serotonergic neurons (Carrera et al., 2008). In the angelshark, hypothalamic serotonergic cells are densely packed at the infundibular ependymal layer (in the inferior lobe of the hypothalamus) and appear to give rise to a number of laterally-projecting fascicles. Hypothalamic and raphe fibers can potentially interact in the distinct plexus overlying some hypothalamic serotonergic cells (Fig. 6E, F) or in more lateral fiber clusters that in coronal sections appear disorganized (Fig. 6C) but may give rise to highly oriented fibers directed toward the habenula (Fig. 6B). In mammals, raphe fibers also enter the hypothalamus; in embryonic rat development, serotonergic axons that reach the infundibular recess (the primordial mammillary complex) are among the first serotonergic fibers to arrive in the forebrain (Lidov and Molliver, 1982; Jacobs and Azmitia, 1992). There are no hypothalamic serotonergic neurons in the rat brain, but comingling between raphe and hypothalamic serotonergic fibers has been shown in monotremes (Manger et al., 2002). Since both raphe and hypothalamic serotonergic neurons are anatomically associated with the ventricular system, one of their original functions may have been to convey CSF-related information to neural tissue (Parent, 1981), thus supporting global functional integration of the CNS.

The observed distribution of serotonergic fibers in the telencephalon allows for a parsimonious description, based on a framework introduced in our recent publications (Janušonis et al., 2020; Janušonis et al., 2023; Mays et al., 2023). Specifically, we assume that serotonergic fibers can be guided by a small set of “deterministic” cues but are inherently “stochastic” in their trajectories.

In the telencephalon, the *deterministic* component can be well captured by the single observation that serotonergic fibers tend to move toward the dorsal pallium (Fig. 5K). The dorsal pallium is an intriguing region in that it may have been absent from the ancestral vertebrate brain and may have emerged several times in different vertebrate lineages, optionally and independently (Striedter and Northcutt, 2020). In particular, in may not be present in amphibians and lungfishes (Striedter and Northcutt, 2020). This hypothesis is consistent with the dual-origin hypothesis of the primate cerebral cortex, a derived dorsal pallial structure with medial and lateral pallial moieties (Pandya et al., 2015). Notably, four distinct fascicles (bands) of serotonergic fibers are present in the angelshark septum (Fig. 5F, I), likely *en route* to the medial side of the dorsal pallium. Such fascicles are also prominent in rodents but there they enter the developing cerebral cortex (the cortical plate) (Lidov and Molliver, 1982; Wallace and Lauder, 1983; Vertes, 1991; Vertes et al., 1999). It suggests that these fascicles, and perhaps the two distinct serotonergic bands that initially “sandwich” the mammalian cortical plate (Wallace and Lauder, 1983; Voigt and de Lima, 1991), are not cortex-specific. In fact, the superficial cortical band, which enters the mammalian cortical marginal zone (Janušonis et al., 2004), has a direct counterpart in the angelshark dorsal pallium (Fig. 5J). We note that the parsimony of this explanation does not rule out the possibility that other regions, such as the habenula, may also attract serotonergic fibers (Fig. 6A, B, D), thus providing additional guidance before the fibers enter the telencephalon. Hypothetically, the attracting cue may a diffusing factor (which, if released by the dorsal pallium, could reach the habenula). These potential gradients have not yet been investigated experimentally or in computer simulations.

The *stochastic* component may be fundamental to the understanding of the apparently complex distribution patterns of serotonergic fibers in different vertebrate species. We have previously shown that the trajectories of serotonergic axons can be modeled as paths of FBM, a continuous-time stochastic process that evolves in space (Janušonis et al., 2020). We also have demonstrated in a supercomputing simulation that simulated FBM-fibers, walking in a geometric shape based on the mouse brain, produces fiber densities that approximate the actual serotonergic fiber densities, with no guiding cues (Janušonis et al., 2023). This result is partly due to a key mathematical property of FBM-fibers, their tendency to accumulate at the shape boundaries (Guggenberger et al., 2019; Vojta et al., 2020). An increased accumulation of serotonergic fibers at the pial and ventricular boundaries has been described by many studies in various brain regions across vertebrate species, but has been interpreted as a neuroanatomical region-specific feature (Martin et al., 1985; Mize and Horner, 1989; Dinopoulos and Dori, 1995; Morin and Meyer-Bernstein, 1999; Bjarkam et al., 2003; 2005; Awasthi et al., 2021; Bhat and Ganesh, 2023). Interestingly, this effect becomes even more pronounced in mice that lack protocadherin-αC2, a cell adhesion protein that is thought to support serotonergic fiber dispersion through self-avoidance (Katori et al., 2009; Katori et al., 2017). Additionally, FBM-trajectories have long-term “memory”: each new step is a random event, but it correlates with the previous steps (history) of the fiber, including its possible reflections at the boundary. Therefore, the three-dimensional geometry of the brain becomes an important factor in the distribution and relative intensity of fiber densities. This conclusion may not appear convincing because brain self-organization and function are typically perceived through the lens of neuroanatomically-defined regions. However, other research shows that brain function may be influenced by brain geometry (Gjorevski et al., 2022; Pang et al., 2023). This change in perspective may uncover fundamental patterns. For example, recent studies have demonstrated strong similarities in neocortical dynamics across mammalian species (Mahon, 2024) and a universal laminar distribution pattern of the power of cortical oscillations across primate neocortical areas (Mendoza-Halliday et al., 2024).

The supercomputing simulation of this study is a significant step in efforts to achieve a deeper understanding of the self-organization of serotonergic fibers. It is the first computational simulation of serotonergic fibers in a non-mammalian brain and it complements our recent supercomputing simulations in the mouse brain (Janušonis et al., 2020; Janušonis et al., 2023). The obtained results show strong similarity to the biological densities of serotonergic fibers and, in particular, highlight the universal tendency of serotonergic fibers to accumulate near tissue boundaries. This simulation does not include deterministic forces (e.g., axon-guiding gradients) which would require a considerable extension of the model, with limited biological information. In an interdisciplinary collaboration, we are currently developing the theory of FBM to incorporate other biologically-relevant elements, such as heterogeneous environments (Wang et al., 2023) or population-induced forces (House et al., 2025).

An interesting possibility exists that some directions chosen by serotonergic fibers are not due to guiding gradients but to the tendency of these fibers to opportunistically travel along major (non-serotonergic) axon tracts. The propensity of serotonergic axons to adhere to many functionally diverse tracts, such as the medial forebrain bundle (Nieuwenhuys et al., 1982) or the fasciculus retroflexus (Beretta et al., 2012), has been noted in early studies and named “epiphytic guidance” (Lidov and Molliver, 1982; Wallace and Lauder, 1983; Jacobs and Azmitia, 1992), in reference to plant epiphytes. Indeed, in shark brains serotonergic fibers could travel along several tracts connecting the diencephalon and telencephalon, such as the tractus pallii or the fasciculus basalis telencephali (Smeets et al., 1983; Smeets, 1998; Hofmann and Northcutt, 2012). Likewise, serotonergic fibers could approach the habenula along the fasciculus retroflexus (Giuliani et al., 2002), as they do in mammalian brains (Jacobs and Azmitia, 1992). In an extreme case, serotonergic fibers could remain entirely stochastic but accumulate around the borders of these tracts, producing an illusion of oriented movement in population-level imaging. However, some individual serotonergic fibers are strongly oriented (Fig. 6B), which suggests that this explanation is insufficient.

Even in the absence of guiding gradients, the simulation produced a strong accumulation of FBM-fibers in the inferior lobe of the hypothalamus (Fig. 10D), the habenula (Fig. 10D), and the dorsal pallium (Fig. 10K), due to their high-curvature contours (Janušonis et al., 2020; Vojta et al., 2020). Some studies have suggested that serotonin itself may stimulate the growth of serotonergic fibers (Azmitia and Whitaker-Azmitia, 1991; Whitaker-Azmitia, 2001). Therefore, these high-density areas can induce a previously absent attracting force, affecting further fiber development. However, this information remains incomplete and sometimes contradictory, with opposite (negative feedback) effects of serotonin also reported (Nishiyama et al., 2002; Kornum et al., 2006; Daubert et al., 2010; Migliarini et al., 2013; Vicenzi et al., 2021; Nazzi et al., 2024).

No detectable border accumulation of serotonergic fibers is present in some angelshark brainstem regions, such as the tectum, perhaps because of frequent reflecting or adhesion events induced by densely packed axon tracts (not necessarily traveling in the same orientation). In theoretical studies, unexpected effects have been observed in particles undergoing FBM motion in heterogeneous, randomly structured environments (Wang et al., 2020). Also, the angelshark cerebellum was almost devoid of serotonergic fibers. This observation is consistent with the low density of serotonergic fibers in the mouse cerebellum (Awasthi et al., 2021). The cerebellum is a highly specialized structure, including the enormous number of granule cells, and may present an otherwise less permissive environment for serotoninergic fibers. It is also may be less physically accessible because of the dense axon tracts connecting it to the medulla (e.g., the cerebellar peduncles in mammals) (Smeets et al., 1983; Smeets, 1998; Montgomery et al., 2012).

Generally, the “flow” of serotonergic fibers in the angelshark forebrain was similar to that of the smallspotted catshark (Carrera et al., 2008). This sequence of fiber progression, in turn, shared considerable similarity with the development of serotonergic axons in the rat brain (Wallace and Lauder, 1983), despite major cytoarchitectonic differences and mammal-specific specializations.

For the purpose of this study, serotonergic fibers were treated as virtually identical in their properties. Recent single-cell RNA-seq studies have revealed an enormous transcriptional diversity of serotonergic neurons (Okaty et al., 2019; Ren et al., 2019; Okaty et al., 2020). These studies represent a major inflection in the field. However, it remains unknown if the same neuron can switch among several transcriptional programs (“attractors”), depending on its environment (Okaty et al., 2019), and whether the current transcriptional program selected by the neuron depends on the path and history of its axon.

In conclusion, we note that the development of many, if not all, axon systems may depend on a dynamic interplay between a mathematically-definable stochastic process and constraining deterministic forces, *at all developmental times* (Fig. 11). Due to the inherent properties of growth cones (Katz et al., 1984; Maskery and Shinbrot, 2005; Betz et al., 2006; Forghani et al., 2023), axons will tend produce stochastic trajectories. Nevertheless, it can result in a rich behavioral repertoire, as axons interact with their physical environment (tissue boundaries, physical obstacles, viscoelastic heterogeneities) and with each other. “Strategically placed” guiding factors (e.g., chemical gradients) are necessary to achieve the genetically predetermined wiring, but they may harness rather than suppress this inherent stochasticity. The depth and richness of such interactions is only beginning to be revealed, with implications for the self-organization, plasticity, and repair of nervous systems.

**Figure 11.**
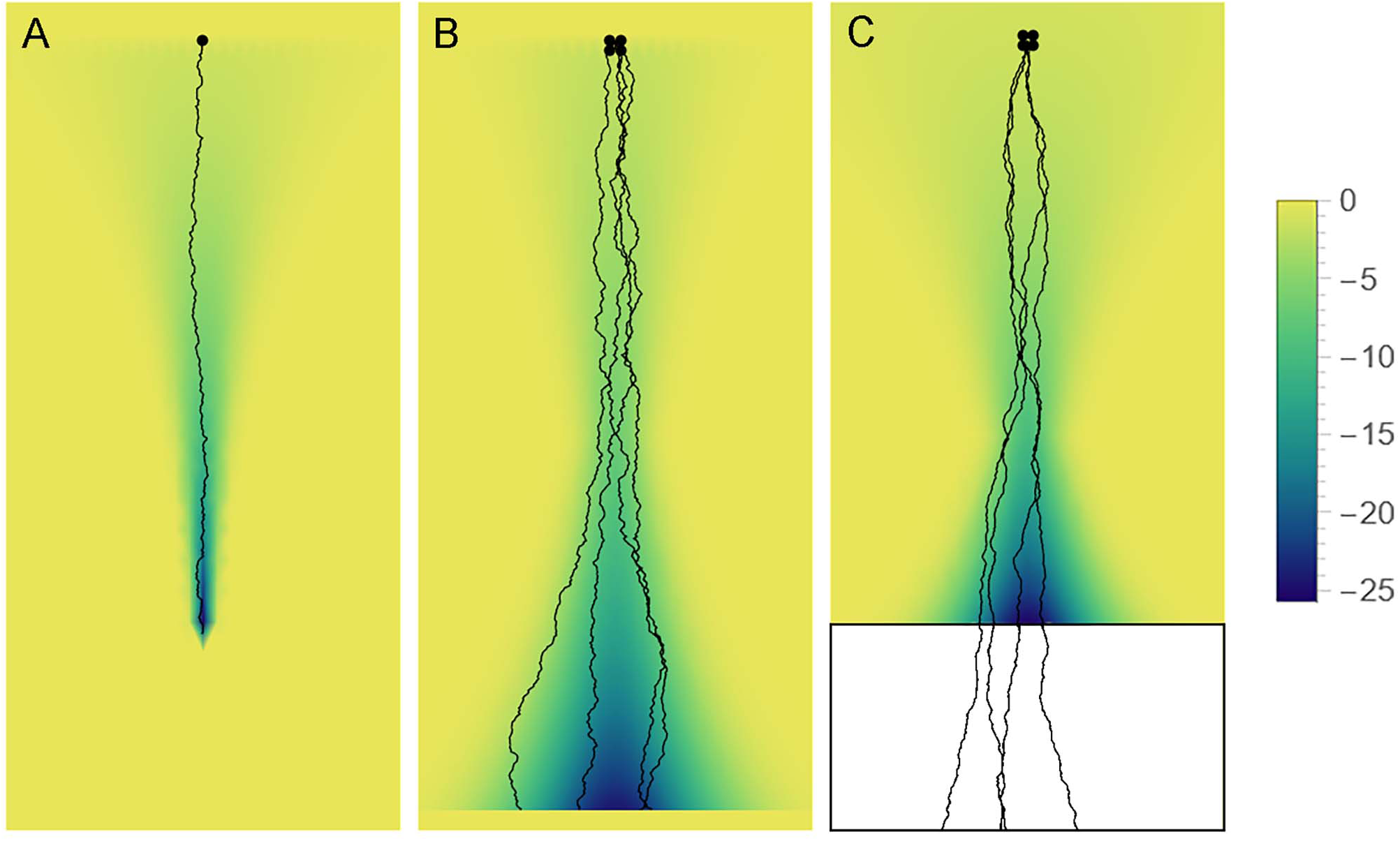
Axon paths may be fundamentally stochastic but can be constrained by “effective potential” landscapes (e.g., axon guiding gradients), depending on the axon type. The colors represent the value of the potential (high = yellow, low = blue). (**A**) Some axons may become trapped in a narrowing potential through and thus be precisely guided to targets (e.g., retinogeniculate axons). (**B**) Some axons may be constrained to more spacious valleys, where their stochasticity may be more apparent (e.g., retinohypothalamic axons). (**C**) Some axons may be guided to specific brain regions in development, to facilitate their dispersal, and then released to produce random walk-like trajectories. Serotonergic axons appear to show some of these properties.

## 6. Author Contributions

SJ designed the project, produced the brain section series, performed all histological and immunohistochemical procedures with imaging, wrote the Mathematica scripts, prepared the figures, conceptualized the biological findings, and wrote the manuscript. TV wrote the FORTRAN scripts and performed all supercomputing simulations. RM contributed to the theory of anomalous diffusion used in supercomputing simulations. SJ, RM, and TV are the Principal Investigators of their respective research programs.

## 8. Acknowledgements

We thank Christoph Pierre (Director of Marine Operations, UCSB) for access to shark specimens. We acknowledge the technical assistance of Dr. Melissa Hingorani, Justin Haiman, Kimia Boldaji, and Caroline Vella (the Janušonis laboratory), and thank Dr. Benjamin Lopez (UCSB Neuroscience Research Institute) for advice in confocal imaging.

## REFERENCES

Aitken, A.R., and Tork, I. (1988). Early development of serotonin-containing neurons and pathways as seen in wholemount preparations of the fetal rat brain. J Comp Neurol 274(1), 32–47. doi: 10.1002/cne.902740105.

Andrews, P.W., Bosyj, C., Brenton, L., Green, L., Gasser, P.J., Lowry, C.A., et al. (2022). All the brain’s a stage for serotonin: the forgotten story of serotonin diffusion across cell membranes. Proc Biol Sci 289(1986), 20221565. doi: 10.1098/rspb.2022.1565.

Antonovaite, N., Hulshof, L.A., Hol, E.M., Wadman, W.J., and Iannuzzi, D. (2021). Viscoelastic mapping of mouse brain tissue: Relation to structure and age. J Mech Behav Biomed Mater 113, 104159. doi: 10.1016/j.jmbbm.2020.104159.

Awasthi, J.R., Tamada, K., Overton, E.T.N., and Takumi, T. (2021). Comprehensive topographical map of the serotonergic fibers in the male mouse brain. J Comp Neurol 529(7), 1391–1429. doi: 10.1002/cne.25027.

Azmitia, E., and Gannon, P. (1983). The ultrastructural localization of serotonin immunoreactivity in myelinated and unmyelinated axons within the medial forebrain bundle of rat and monkey. J Neurosci 3(10), 2083–2090. doi: 10.1523/jneurosci.03-10-02083.1983.

Azmitia, E.C., and Whitaker-Azmitia, P.M. (1991). Awakening the sleeping giant: anatomy and plasticity of the brain serotonergic system. J Clin Psychiatry 52 Suppl, 4–16.

Barca, A., Vacca, F., Vizioli, J., Drago, F., Vetrugno, C., Verri, T., et al. (2017). Molecular and expression analysis of the allograft inflammatory factor 1 (AIF-1) in the coelomocytes of the common sea urchin *Paracentrotus lividus*. Fish Shellfish Immunol 71, 136–143. doi: 10.1016/j.fsi.2017.09.078.

Beretta, C.A., Dross, N., Guiterrez-Triana, J.A., Ryu, S., and Carl, M. (2012). Habenula circuit development: past, present, and future. Front Neurosci 6, 51. doi: 10.3389/fnins.2012.00051.

Betz, T., Lim, D., and Käs, J.A. (2006). Neuronal growth: a bistable stochastic process. Phys Rev Lett 96(9), 098103. doi: 10.1103/PhysRevLett.96.098103.

Bhat, S.K., and Ganesh, C.B. (2023). Organization of serotonergic system in *Sphaerotheca breviceps* (Dicroglossidae) tadpole brain. Cell Tissue Res 391(1), 67–86. doi: 10.1007/s00441-022-03709-7.

Biagini, F., Hu, Y., Oksendal, B., and Zhang, T. (2010). Stochastic Calculus for Fractional Brownian Motion and Applications. London: Springer.

Biradar, A., and Ganesh, C.B. (2024). Serotonin-immunoreactivity in the brain of the cichlid fish *Oreochromis mossambicus*. Anat Rec (Hoboken) 307(2), 320–344. doi: 10.1002/ar.25204.

Bjarkam, C.R., Sørensen, J.C., and Geneser, F.A. (2003). Distribution and morphology of serotonin-immunoreactive axons in the hippocampal region of the New Zealand white rabbit. I. Area dentata and hippocampus. Hippocampus 13(1), 21–37. doi: 10.1002/hipo.10042.

Bjarkam, C.R., Sørensen, J.C., and Geneser, F.A. (2005). Distribution and morphology of serotonin-immunoreactive axons in the retrohippocampal areas of the New Zealand white rabbit. Anat Embryol (Berl) 210(3), 199–207. doi: 10.1007/s00429-005-0004-x.

Blumenfeld, H. (2022). Neuroanatomy Through Clinical Cases. New York: Oxford University Press.

Bockaert, J., Claeysen, S., Bécamel, C., Dumuis, A., and Marin, P. (2006). Neuronal 5-HT metabotropic receptors: fine-tuning of their structure, signaling, and roles in synaptic modulation. Cell Tissue Res 326(2), 553–572. doi: 10.1007/s00441-006-0286-1.

Butler, A.B., and Hodos, W. (2005). Comparative Vertebrate Neuroanatomy (2nd edition). Hoboken: Wiley-Interscience.

Carrera, I., Molist, P., Anadon, R., and Rodriguez-Moldes, I. (2008). Development of the serotoninergic system in the central nervous system of a shark, the lesser spotted dogfish *Scyliorhinus canicula*. J Comp Neurol 511(6), 804–831. doi: 10.1002/cne.21857.

Cazettes, F., Reato, D., Morais, J.P., Renart, A., and Mainen, Z.F. (2021). Phasic activation of dorsal raphe serotonergic neurons increases pupil size. Curr Biol 31(1), 192–197.e194. doi: 10.1016/j.cub.2020.09.090.

Chaudhuri, O., Cooper-White, J., Janmey, P.A., Mooney, D.J., and Shenoy, V.B. (2020). Effects of extracellular matrix viscoelasticity on cellular behaviour. Nature 584(7822), 535–546. doi: 10.1038/s41586-020-2612-2.

Chen, W.V., Nwakeze, C.L., Denny, C.A., O’Keeffe, S., Rieger, M.A., Mountoufaris, G., et al. (2017). Pcdhαc2 is required for axonal tiling and assembly of serotonergic circuitries in mice. Science 356(6336), 406–411. doi: 10.1126/science.aal3231.

Cheung, A., Konno, K., Imamura, Y., Matsui, A., Abe, M., Sakimura, K., et al. (2023). Neurexins in serotonergic neurons regulate neuronal survival, serotonin transmission, and complex mouse behaviors. Elife 12. doi: 10.7554/eLife.85058.

Compagno, L., Dando, M., and Sarah, F. (2005). Sharks of the World. Princeton: Princeton University Press.

Cooke, P., Janowitz, H., and Dougherty, S.E. (2022). Neuronal redevelopment and the regeneration of neuromodulatory axons in the adult mammalian central nervous system. Front Cell Neurosci 16, 872501. doi: 10.3389/fncel.2022.872501.

Daubert, E.A., Heffron, D.S., Mandell, J.W., and Condron, B.G. (2010). Serotonergic dystrophy induced by excess serotonin. Mol Cell Neurosci 44(3), 297–306. doi: 10.1016/j.mcn.2010.04.001.

de Bellard, M.E. (2016). Myelin in cartilaginous fish. Brain Res 1641(Pt A), 34–42. doi: 10.1016/j.brainres.2016.01.013.

Dey, S., Surendran, D., Engberg, O., Gupta, A., Fanibunda, S.E., Das, A., et al. (2021). Altered membrane mechanics provides a receptor-independent pathway for serotonin action. Chemistry 27(27), 7533–7541. doi: 10.1002/chem.202100328.

Dinopoulos, A., and Dori, I. (1995). The development of the serotonergic fiber network of the lateral ventricles of the rat brain: a light and electron microscopic immunocytochemical analysis. Exp Neurol 133(1), 73–84. doi: 10.1006/exnr.1995.1009.

Dohare, S., Hernandez-Garcia, J.F., Lan, Q., Rahman, P., Mahmood, A.R., and Sutton, R.S. (2024). Loss of plasticity in deep continual learning. Nature 632(8026), 768–774. doi: 10.1038/s41586-024-07711-7.

Donovan, L.J., Spencer, W.C., Kitt, M.M., Eastman, B.A., Lobur, K.J., Jiao, K., et al. (2019). Lmx1b is required at multiple stages to build expansive serotonergic axon architectures. Elife 8, e48788. doi: 10.7554/eLife.48788.

Ebbesson, L.O., Holmqvist, B., Ostholm, T., and Ekström, P. (1992). Transient serotonin-immunoreactive neurons coincide with a critical period of neural development in coho salmon (*Oncorhynchus kisutch*). Cell Tissue Res 268(2), 389–392. doi: 10.1007/bf00318807.

Flood, Z.C., Engel, D.L., Simon, C.C., Negherbon, K.R., Murphy, L.J., Tamavimok, W., et al. (2012). Brain growth trajectories in mouse strains with central and peripheral serotonin differences: relevance to autism models. Neuroscience 210, 286–295. doi: 10.1016/j.neuroscience.2012.03.010.

Fonseca, E.M., Noronha-de-Souza, C.R., Bícego, K.C., Branco, L.G.S., and Gargaglioni, L.H. (2021). 5-HT neurons of the medullary raphe contribute to respiratory control in toads. Respir Physiol Neurobiol 293, 103717. doi: 10.1016/j.resp.2021.103717.

Forghani, R., Chandrasekaran, A., Papoian, G., and Giniger, E. (2023). A new view of axon growth and guidance grounded in the stochastic dynamics of actin networks. Open Biol 13(6), 220359. doi: 10.1098/rsob.220359.

Fujita, T., Aoki, N., Mori, C., Homma, K.J., and Yamaguchi, S. (2023). Molecular biology of serotonergic systems in avian brains. Front Mol Neurosci 16, 1226645. doi: 10.3389/fnmol.2023.1226645.

Furness, J.B., and Stebbing, M.J. (2018). The first brain: Species comparisons and evolutionary implications for the enteric and central nervous systems. Neurogastroenterol Motil 30(2). doi: 10.1111/nmo.13234.

Gershon, M.D., and Margolis, K.G. (2021). The gut, its microbiome, and the brain: connections and communications. J Clin Invest 131(18). doi: 10.1172/jci143768.

Gianni, G., and Pasqualetti, M. (2023). Wiring and volume transmission: An overview of the dual modality for serotonin neurotransmission. ACS Chem Neurosci 14(23), 4093–4104. doi: 10.1021/acschemneuro.3c00648.

Giuliani, A., Minelli, D., Quaglia, A., and Villani, L. (2002). Telencephalo-habenulo-interpeduncular connections in the brain of the shark *Chiloscyllium arabicum*. Brain Res 926(1-2), 186–190. doi: 10.1016/s0006-8993(01)03310-8.

Gjorevski, N., Nikolaev, M., Brown, T.E., Mitrofanova, O., Brandenberg, N., DelRio, F.W., et al. (2022). Tissue geometry drives deterministic organoid patterning. Science 375(6576), eaaw9021. doi: 10.1126/science.aaw9021.

Guggenberger, T., Pagnini, G., Vojta, T., and Metzler, R. (2019). Fractional Brownian motion in a finite interval: correlations effect depletion or accretion zones of particles near boundaries. New J. Phys. 21(022002).

Hawthorne, A.L., Hu, H., Kundu, B., Steinmetz, M.P., Wylie, C.J., Deneris, E.S., et al. (2011). The unusual response of serotonergic neurons after CNS Injury: lack of axonal dieback and enhanced sprouting within the inhibitory environment of the glial scar. J Neurosci 31(15), 5605–5616. doi: 10.1523/jneurosci.6663-10.2011.

Hingorani, M., Viviani, A.M.L., Sanfilippo, J.E., and Janušonis, S. (2022). High-resolution spatiotemporal analysis of single serotonergic axons in an in vitro system. Front Neurosci 16, 994735. doi: 10.3389/fnins.2022.994735.

Hofmann, M.H., and Northcutt, R.G. (2012). Forebrain organization in elasmobranchs. Brain Behav Evol 80(2), 142–151. doi: 10.1159/000339874.

Hornung, J.P. (2003). The human raphe nuclei and the serotonergic system. J Chem Neuroanat 26(4), 331–343. doi: 10.1016/j.jchemneu.2003.10.002.

House, J., Janušonis, S., Metzler, R., and Vojta, T. (2025). Fractional Brownian motion with mean-density interaction. arXiv 2503.15255v1 doi: 10.48550/arXiv.2503.15255.

Jacobs, B.L., and Azmitia, E.C. (1992). Structure and function of the brain serotonin system. Physiol Rev 72(1), 165–229. doi: 10.1152/physrev.1992.72.1.165.

Janušonis, S. (2018). Some galeomorph sharks express a mammalian microglia-specific protein in radial ependymoglia of the telencephalon. Brain Behav Evol 91(1), 17–30. doi: 10.1159/000484196.

Janušonis, S., and Detering, N. (2019). A stochastic approach to serotonergic fibers in mental disorders. Biochimie 161, 15–22. doi: 10.1016/j.biochi.2018.07.014.

Janušonis, S., Detering, N., Metzler, R., and Vojta, T. (2020). Serotonergic axons as fractional Brownian motion paths: Insights into the self-organization of regional densities. Front Comput Neurosci 14, 56. doi: 10.3389/fncom.2020.00056.

Janušonis, S., Gluncic, V., and Rakic, P. (2004). Early serotonergic projections to Cajal-Retzius cells: relevance for cortical development. J Neurosci 24(7), 1652–1659. doi: 10.1523/jneurosci.4651-03.2004.

Janušonis, S., Haiman, J.H., Metzler, R., and Vojta, T. (2023). Predicting the distribution of serotonergic axons: a supercomputing simulation of reflected fractional Brownian motion in a 3D-mouse brain model. Front Comput Neurosci 17, 1189853. doi: 10.3389/fncom.2023.1189853.

Jin, Y., Dougherty, S.E., Wood, K., Sun, L., Cudmore, R.H., Abdalla, A., et al. (2016). Regrowth of serotonin axons in the adult mouse brain following injury. Neuron 91(4), 748–762. doi: 10.1016/j.neuron.2016.07.024.

Jordan, J.E.L., Bertalan, G., Meyer, T., Tzschätzsch, H., Gauert, A., Bramè, L., et al. (2022). Microscopic multifrequency MR elastography for mapping viscoelasticity in zebrafish. Magn Reson Med 87(3), 1435–1445. doi: 10.1002/mrm.29066.

Kajstura, T.J., Dougherty, S.E., and Linden, D.J. (2018). Serotonin axons in the neocortex of the adult female mouse regrow after traumatic brain injury. J Neurosci Res 96(4), 512–526. doi: 10.1002/jnr.24059.

Katori, S., Hamada, S., Noguchi, Y., Fukuda, E., Yamamoto, T., Yamamoto, H., et al. (2009). Protocadherin-alpha family is required for serotonergic projections to appropriately innervate target brain areas. J Neurosci 29(29), 9137–9147. doi: 10.1523/jneurosci.5478-08.2009.

Katori, S., Noguchi-Katori, Y., Okayama, A., Kawamura, Y., Luo, W., Sakimura, K., et al. (2017). Protocadherin-αC2 is required for diffuse projections of serotonergic axons. Sci Rep 7(1), 15908. doi: 10.1038/s41598-017-16120-y.

Katz, M.J., George, E.B., and Gilbert, L.J. (1984). Axonal elongation as a stochastic walk. Cell Motil 4(5), 351–370. doi: 10.1002/cm.970040505.

Kitt, M.M., Tabuchi, N., Spencer, W.C., Robinson, H.L., Zhang, X.L., Eastman, B.A., et al. (2022). An adult-stage transcriptional program for survival of serotonergic connectivity. Cell Rep 39(3), 110711. doi: 10.1016/j.celrep.2022.110711.

Kiyasova, V., and Gaspar, P. (2011). Development of raphe serotonin neurons from specification to guidance. Eur J Neurosci 34(10), 1553–1562. doi: 10.1111/j.1460-9568.2011.07910.x.

Kornum, B.R., Licht, C.L., Weikop, P., Knudsen, G.M., and Aznar, S. (2006). Central serotonin depletion affects rat brain areas differently: a qualitative and quantitative comparison between different treatment schemes. Neurosci Lett 392(1-2), 129–134. doi: 10.1016/j.neulet.2005.09.013.

Lee, C., Zhang, Z., and Janušonis, S. (2022). Brain serotonergic fibers suggest anomalous diffusion-based dropout in artificial neural networks. Front Neurosci 16, 949934. doi: 10.3389/fnins.2022.949934.

Lesch, K.P., and Waider, J. (2012). Serotonin in the modulation of neural plasticity and networks: implications for neurodevelopmental disorders. Neuron 76(1), 175–191. doi: 10.1016/j.neuron.2012.09.013.

Lidov, H.G., and Molliver, M.E. (1982). An immunohistochemical study of serotonin neuron development in the rat: ascending pathways and terminal fields. Brain Res Bull 8(4), 389–430. doi: 10.1016/0361-9230(82)90077-6.

Lillesaar, C. (2011). The serotonergic system in fish. J Chem Neuroanat 41(4), 294–308. doi: 10.1016/j.jchemneu.2011.05.009.

López-Romero, F.A., Stumpf, S., Pfaff, C., Marramà, G., Johanson, Z., and Kriwet, J. (2020). Evolutionary trends of the conserved neurocranium shape in angel sharks (Squatiniformes, Elasmobranchii). Sci Rep 10(1), 12582. doi: 10.1038/s41598-020-69525-7.

MacDonnell, M.F. (1980). Cerebrospinal fluid contacting and supraependymal mesencephalic trigeminal cells in the blue and mako sharks. A scanning electron microscopic study. Brain Behav Evol 17(2), 164–177. doi: 10.1159/000121796.

MacDonnell, M.F. (1983). Mesencephalic trigeminal midline ridge formation in sharks, a proposed circumventricular organ: developmental aspects. Anat Rec 206(3), 319–327. doi: 10.1002/ar.1092060311.

MacDonnell, M.F. (1984). Circumventricular mesencephalic trigeminal midline ridge formation in cartilaginous fishes: species variations. Brain Behav Evol 24(2-3), 124–134. doi: 10.1159/000121310.

MacDonnell, M.F. (1989). Sub/supraependymal axonal net in the brains of sharks and probable targets in parasynaptic relationship. Brain Behav Evol 34(4), 201–211. doi: 10.1159/000116506.

Mahon, S. (2024). Variation and convergence in the morpho-functional properties of the mammalian neocortex. Front Syst Neurosci 18, 1413780. doi: 10.3389/fnsys.2024.1413780.

Maisey, J.G., Ehret, D.J., and Denton, J.S.S. (2020). A new genus of late Cretaceous angel shark (Elasmobranchii; Squatinidae), with comments on squatinid phylogeny. American Museum Novitates 3954, 1–29. doi: 10.1206/3954.1.

Makse, H.A., Havlin, S., Schwartz, M., and Stanley, H.E. (1996). Method for generating long-range correlations for large systems. Physical Review E 53, 5445–5449.

Mamounas, L.A., Blue, M.E., Siuciak, J.A., and Altar, C.A. (1995). Brain-derived neurotrophic factor promotes the survival and sprouting of serotonergic axons in rat brain. J Neurosci 15(12), 7929–7939.

Mandelbrot, B.B., and Van Ness, J.W. (1968). Fractional Brownian motions, fractional noises and applications. SIAM Review 10, 422–437.

Manger, P.R., Fahringer, H.M., Pettigrew, J.D., and Siegel, J.M. (2002). The distribution and morphological characteristics of serotonergic cells in the brain of monotremes. Brain Behav Evol 60(5), 315–332. doi: 10.1159/000067194.

Martin, G.F., DeLorenzo, G., Ho, R.H., Humbertson, A.O., Jr., and Waltzer, R. (1985). Serotonergic innervation of the forebrain in the North American opossum. Brain Behav Evol 26(3-4), 196–228. doi: 10.1159/000118776.

Maskery, S., and Shinbrot, T. (2005). Deterministic and stochastic elements of axonal guidance. Annu Rev Biomed Eng 7, 187–221. doi: 10.1146/annurev.bioeng.7.060804.100446.

Mays, K.C., Haiman, J.H., and Janušonis, S. (2023). An experimental platform for stochastic analyses of single serotonergic fibers in the mouse brain. Front Neurosci 17, 1241919. doi: 10.3389/fnins.2023.1241919.

McCormick, H.W., Allen, T., and Young, W.E. (1963). Shadows in the Sea: The Sharks, Skates and Rays. New York: Weathervane Books.

Mendoza-Halliday, D., Major, A.J., Lee, N., Lichtenfeld, M.J., Carlson, B., Mitchell, B., et al. (2024). A ubiquitous spectrolaminar motif of local field potential power across the primate cortex. Nat Neurosci 27(3), 547–560. doi: 10.1038/s41593-023-01554-7.

Migliarini, S., Pacini, G., Pelosi, B., Lunardi, G., and Pasqualetti, M. (2013). Lack of brain serotonin affects postnatal development and serotonergic neuronal circuitry formation. Mol Psychiatry 18(10), 1106–1118. doi: 10.1038/mp.2012.128.

Mize, R.R., and Horner, L.H. (1989). Origin, distribution, and morphology of serotonergic afferents to the cat superior colliculus: a light and electron microscope immunocytochemistry study. Exp Brain Res 75(1), 83–98. doi: 10.1007/bf00248533.

Montalbano, A., Waider, J., Barbieri, M., Baytas, O., Lesch, K.P., Corradetti, R., et al. (2015). Cellular resilience: 5-HT neurons in Tph2(-/-) mice retain normal firing behavior despite the lack of brain 5-HT. Eur Neuropsychopharmacol 25(11), 2022–2035. doi: 10.1016/j.euroneuro.2015.08.021.

Montgomery, J.C., Bodznick, D., and Yopak, K.E. (2012). The cerebellum and cerebellum-like structures of cartilaginous fishes. Brain Behav Evol 80(2), 152–165. doi: 10.1159/000339868.

Morin, L.P., and Meyer-Bernstein, E.L. (1999). The ascending serotonergic system in the hamster: comparison with projections of the dorsal and median raphe nuclei. Neuroscience 91(1), 81–105. doi: 10.1016/s0306-4522(98)00585-5.

Mosienko, V., Beis, D., Pasqualetti, M., Waider, J., Matthes, S., Qadri, F., et al. (2015). Life without brain serotonin: reevaluation of serotonin function with mice deficient in brain serotonin synthesis. Behav Brain Res 277, 78–88. doi: 10.1016/j.bbr.2014.06.005.

Nazzi, S., Maddaloni, G., Pratelli, M., and Pasqualetti, M. (2019). Fluoxetine induces morphological rearrangements of serotonergic fibers in the hippocampus. ACS Chem Neurosci 10(7), 3218–3224. doi: 10.1021/acschemneuro.8b00655.

Nazzi, S., Picchi, M., Migliarini, S., Maddaloni, G., Barsotti, N., and Pasqualetti, M. (2024). Reversible morphological remodeling of prefrontal and hippocampal serotonergic fibers by Fluoxetine. ACS Chem Neurosci 15(8), 1702–1711. doi: 10.1021/acschemneuro.3c00837.

New, J.G. (2001). Comparative neurobiology of the elasmobranch cerebellum: theme and variations on a sensorimotor interface. Environmental Biology of Fishes 60, 93–108. doi: 10.1023/A:1007631405904.

Nieuwenhuys, R., Geeraedts, L.M., and Veening, J.G. (1982). The medial forebrain bundle of the rat. I. General introduction. J Comp Neurol 206(1), 49–81. doi: 10.1002/cne.902060106.

Nishiyama, H., Takemura, M., Takeda, T., and Itohara, S. (2002). Normal development of serotonergic neurons in mice lacking S100β. Neurosci Lett 321(1-2), 49–52. doi: 10.1016/s0304-3940(01)02549-6.

Ogelman, R., Gomez Wulschner, L.E., Hoelscher, V.M., Hwang, I.W., Chang, V.N., and Oh, W.C. (2024). Serotonin modulates excitatory synapse maturation in the developing prefrontal cortex. Nat Commun 15(1), 1368. doi: 10.1038/s41467-024-45734-w.

Okaty, B.W., Commons, K.G., and Dymecki, S.M. (2019). Embracing diversity in the 5-HT neuronal system. Nat Rev Neurosci 20(7), 397–424. doi: 10.1038/s41583-019-0151-3.

Okaty, B.W., Sturrock, N., Escobedo Lozoya, Y., Chang, Y., Senft, R.A., Lyon, K.A., et al. (2020). A single-cell transcriptomic and anatomic atlas of mouse dorsal raphe Pet1 neurons. Elife 9. doi: 10.7554/eLife.55523.

Page, C.E., Epperson, C.N., Novick, A.M., Duffy, K.A., and Thompson, S.M. (2024). Beyond the serotonin deficit hypothesis: communicating a neuroplasticity framework of major depressive disorder. Mol Psychiatry 29(12), 3802–3813. doi: 10.1038/s41380-024-02625-2.

Pandya, D.N., Seltzer, B., Petrides, M., and Cipolloni, P.B. (2015). Cerebral Cortex: Architecture, Connections, and the Dual Origin Concept. Oxford University Press.

Pang, J.C., Aquino, K.M., Oldehinkel, M., Robinson, P.A., Fulcher, B.D., Breakspear, M., et al. (2023). Geometric constraints on human brain function. Nature 618(7965), 566–574. doi: 10.1038/s41586-023-06098-1.

Papadopoulos, G.C., Parnavelas, J.G., and Buijs, R. (1987). Monoaminergic fibers form conventional synapses in the cerebral cortex. Neurosci Lett 76(3), 275–279. doi: 10.1016/0304-3940(87)90414-9.

Parent, A. (1981). Comparative anatomy of the serotoninergic systems. J Physiol (Paris) 77(2-3), 147–156.

Paxinos, G., Kassem, M.S., Kirkcaldie, M., and P., C. (2022). Chemoarchitectonic Atlas of the Rat Brain. London: Academic Press.

Qian, H. (2003). Fractional Brownian motion and fractional Gaussian noise, in Processes with Long-Range Correlations: Theory and Applications, eds. G. Rangarajan & M. Ding. (Berlin/Heidelberg: Springer), 22-33.

Ren, J., Friedmann, D., Xiong, J., Liu, C.D., Ferguson, B.R., Weerakkody, T., et al. (2018). Anatomically defined and functionally distinct dorsal raphe serotonin sub-systems. Cell 175(2), 472–487.e420. doi: 10.1016/j.cell.2018.07.043.

Ren, J., Isakova, A., Friedmann, D., Zeng, J., Grutzner, S.M., Pun, A., et al. (2019). Single-cell transcriptomes and whole-brain projections of serotonin neurons in the mouse dorsal and median raphe nuclei. Elife 8. doi: 10.7554/eLife.49424.

Rizzo, D.J., White, J.D., Spedden, E., Wiens, M.R., Kaplan, D.L., Atherton, T.J., et al. (2013). Neuronal growth as diffusion in an effective potential. Phys Rev E Stat Nonlin Soft Matter Phys 88(4), 042707. doi: 10.1103/PhysRevE.88.042707.

Roberts, B.L., and Witkovsky, P. (1975). A functional analysis of the mesencephalic nucleus of the fifth nerve in the selachian brain. Proc R Soc Lond B Biol Sci 190(1101), 473–495. doi: 10.1098/rspb.1975.0107.

Rodrigues, S.L., Maseko, B.C., Ihunwo, A.O., Fuxe, K., and Manger, P.R. (2008). Nuclear organization and morphology of serotonergic neurons in the brain of the Nile crocodile, *Crocodylus niloticus*. J Chem Neuroanat 35(1), 133–145. doi: 10.1016/j.jchemneu.2007.08.007.

Santos, T.E., Schaffran, B., Broguière, N., Meyn, L., Zenobi-Wong, M., and Bradke, F. (2020). Axon growth of CNS neurons in three dimensions is amoeboid and independent of adhesions. Cell Rep 32(3), 107907. doi: 10.1016/j.celrep.2020.107907.

Shimada, T., Kohyama, K., Yoshida, T., and Yamagata, K. (2024). Neuritin controls axonal branching in serotonin neurons: A possible mediator involved in the regulation of depressive and anxiety behaviors via FGF signaling. J Neurosci 44(41). doi: 10.1523/jneurosci.0129-23.2024.

Slaten, E.R., Hernandez, M.C., Albay, R., 3rd, Lavian, R., and Janušonis, S. (2010). Transient expression of serotonin 5-HT_4_ receptors in the mouse developing thalamocortical projections. Dev Neurobiol 70(3), 165–181. doi: 10.1002/dneu.20775.

Smeets, W.J.A.J. (1998). Cartilaginous Fishes, in The Central Nervous System of Vertebrates, eds. R. Nieuwenhuys, H.J. Ten Donkelaar & C. Nicholson. (Berlin: Springer-Verlag).

Smeets, W.J.A.J., Nieuwenhuys, R., and Roberts, B.L. (1983). The Central Nervous System of Cartilaginous Fishes: Structure and Functional Correlations. New York: Springer-Verlag.

Smiley, J.F., and Goldman-Rakic, P.S. (1996). Serotonergic axons in monkey prefrontal cerebral cortex synapse predominantly on interneurons as demonstrated by serial section electron microscopy. J Comp Neurol 367(3), 431–443. doi: 10.1002/(sici)1096-9861(19960408)367:3<431::Aid-cne8>3.0.Co;2-6.

Striedter, G.F., and Northcutt, R.G. (2020). Brains Through Time. New York: Oxford University Press.

Stuesse, S.L., Cruce, W.L., and Northcutt, R.G. (1991). Localization of serotonin, tyrosine hydroxylase, and leu-enkephalin immunoreactive cells in the brainstem of the horn shark, *Heterodontus francisci*. J Comp Neurol 308(2), 277–292. doi: 10.1002/cne.903080211.

Stuesse, S.L., Stuesse, D.C., and Cruce, W.L. (1995). Raphe nuclei in three cartilaginous fishes, *Hydrolagus colliei*, *Heterodontus francisci*, and *Squalus acanthias*. J Comp Neurol 358(3), 414–427. doi: 10.1002/cne.903580308.

Teissier, A., Soiza-Reilly, M., and Gaspar, P. (2017). Refining the role of 5-HT in postnatal development of brain circuits. Front Cell Neurosci 11, 139. doi: 10.3389/fncel.2017.00139.

Verkhratsky, A., Arranz, A.M., Ciuba, K., and Pękowska, A. (2022). Evolution of neuroglia. Ann N Y Acad Sci 1518(1), 120–130. doi: 10.1111/nyas.14917.

Vertes, R.P. (1991). A PHA-L analysis of ascending projections of the dorsal raphe nucleus in the rat. J Comp Neurol 313(4), 643–668. doi: 10.1002/cne.903130409.

Vertes, R.P., Fortin, W.J., and Crane, A.M. (1999). Projections of the median raphe nucleus in the rat. J Comp Neurol 407(4), 555–582.

Vicenzi, S., Foa, L., and Gasperini, R.J. (2021). Serotonin functions as a bidirectional guidance molecule regulating growth cone motility. Cell Mol Life Sci 78(5), 2247–2262. doi: 10.1007/s00018-020-03628-2.

Voigt, T., and de Lima, A.D. (1991). Serotoninergic innervation of the ferret cerebral cortex. II. Postnatal development. J Comp Neurol 314(2), 415–428. doi: 10.1002/cne.903140215.

Vojta, T., Halladay, S., Skinner, S., Janušonis, S., Guggenberger, T., and Metzler, R. (2020). Reflected fractional Brownian motion in one and higher dimensions. Phys Rev E 102(3-1), 032108. doi: 10.1103/PhysRevE.102.032108.

Wada, A.H.O., and Vojta, T. (2018). Fractional Brownian motion with a reflecting wall. Phys Rev E 97(2-1), 020102. doi: 10.1103/PhysRevE.97.020102.

Wake, H., Moorhouse, A.J., Jinno, S., Kohsaka, S., and Nabekura, J. (2009). Resting microglia directly monitor the functional state of synapses in vivo and determine the fate of ischemic terminals. J Neurosci 29(13), 3974–3980. doi: 10.1523/jneurosci.4363-08.2009.

Wallace, J.A., and Lauder, J.M. (1983). Development of the serotonergic system in the rat embryo: an immunocytochemical study. Brain Res Bull 10(4), 459–479. doi: 10.1016/0361-9230(83)90144-2.

Wang, W., Balcerek, M., Burnecki, K., Chechkin, A.V., Janušonis, S., Ślęzak, J., et al. (2023). Memory-multi-fractional Brownian motion with continuous correlations. Phys. Rev. Res. 5, L032025. doi: 10.1103/PhysRevResearch.5.L032025.

Wang, W., Seno, F., Sokolov, I.M., Chechkin, A.V., and Metzler, R. (2020). Unexpected crossovers in correlated random-diffusivity processes. New Journal of Physics 22, 083041. doi: 10.1088/1367-2630/aba390.

Westlund, K.N., Lu, Y., Coggeshall, R.E., and Willis, W.D. (1992). Serotonin is found in myelinated axons of the dorsolateral funiculus in monkeys. Neurosci Lett 141(1), 35–38. doi: 10.1016/0304-3940(92)90328-5.

Whitaker-Azmitia, P.M. (2001). Serotonin and brain development: role in human developmental diseases. Brain Res Bull 56(5), 479–485. doi: 10.1016/s0361-9230(01)00615-3.

Witkovsky, P., and Roberts, B.L. (1975). The light microscopical structure of the mesencephalic nucleus of the fifth nerve in the selachian brain. Proc R Soc Lond B Biol Sci 190(1101), 457–471. doi: 10.1098/rspb.1975.0106.

Witkovsky, P., and Roberts, B.L. (1976). Electron microscopic observations of the mesencephalic nucleus of the fifth nerve in the Selachian brain. J Neurocytol 5(6), 643–660. doi: 10.1007/bf01181578.

Yip, P.K., Schmitzberger, M., Al-Hasan, M., George, J., Tripoliti, E., Michael-Titus, A.T., et al. (2020). Serotonin expression in the song circuitry of adult male zebra finches. Neuroscience 444, 170–182. doi: 10.1016/j.neuroscience.2020.06.018.

Yopak, K.E. (2012). Neuroecology of cartilaginous fishes: the functional implications of brain scaling. J Fish Biol 80(5), 1968–2023. doi: 10.1111/j.1095-8649.2012.03254.x.

Yopak, K.E., McMeans, B.C., Mull, C.G., Feindel, K.W., Kovacs, K.M., Lydersen, C., et al. (2019). Comparative brain morphology of the greenland and pacific sleeper sharks and its functional implications. Sci Rep 9(1), 10022. doi: 10.1038/s41598-019-46225-5.

Zalc, B. (2016). The acquisition of myelin: An evolutionary perspective. Brain Res 1641(Pt A), 4–10. doi: 10.1016/j.brainres.2015.09.005.

